# A bivalent self-amplifying RNA vaccine against yellow fever and Zika viruses

**DOI:** 10.1101/2025.01.31.635934

**Authors:** Peter Battisti, Matthew R. Ykema, Darshan N. Kasal, Madeleine F. Jennewein, Samuel Beaver, Abbie E. Weight, Derek Hanson, Jasneet Singh, Julie Bakken, Noah Cross, Pauline Fusco, Jacob Archer, Sierra Reed, Alana Gerhardt, Justin G. Julander, Corey Casper, Emily A. Voigt

## Abstract

**Introduction:** Yellow fever (YFV) and Zika (ZIKV) viruses cause significant morbidity and mortality, despite the existence of an approved YFV vaccine and the development of multiple ZIKV vaccine candidates to date. New technologies may improve access to vaccines against these pathogens. We previously described a nanostructured lipid carrier (NLC)-delivered self-amplifying RNA (saRNA) vaccine platform with excellent thermostability and immunogenicity, appropriate for prevention of tropical infectious diseases.

**Methods:** YFV and ZIKV prM-E antigen-expressing saRNA constructs were created using a TC-83 strain Venezuelan equine encephalitis virus-based replicon and complexed with NLC by simple mixing. Monovalent and bivalent vaccine formulations were injected intramuscularly into C57BL/6 mice and Syrian golden hamsters, and the magnitude, durability, and protective efficacy of the resulting immune responses were then characterized.

**Results and discussion:** Monovalent vaccines established durable neutralizing antibody responses to their respective flaviviral targets, with little evidence of cross-neutralization. Both vaccines additionally elicited robust antigen-reactive CD4^+^ and CD8^+^ T cell populations. Notably, humoral responses to YFV saRNA-NLC vaccination were comparable to those in YF-17D-vaccinated animals. Bivalent formulations established humoral and cellular responses against both viral targets, commensurate to those established by monovalent vaccines, without evidence of saRNA interference or immune competition. Finally, both monovalent and bivalent vaccines completely protected mice and hamsters against lethal ZIKV and YFV challenge. We present a bivalent saRNA-NLC vaccine against YFV and ZIKV capable of inducing robust and efficacious neutralizing antibody and cellular immune responses against both viruses. These data support the development of other multivalent saRNA-based vaccines against infectious diseases.

## INTRODUCTION

Yellow fever virus (YFV) and Zika virus (ZIKV) are mosquito-borne flaviviruses transmitted by *Aedes* species mosquitos, which cause infectious disease with a significant impact on global health. YFV is found in tropical regions of Africa and South America, and causes fever, jaundice, and liver damage that may become fatal.(1) The live-attenuated YF-17D vaccine against YFV has been a gold standard vaccine since its introduction in 1937. The YF-17D vaccine is highly effective in disease prevention, with a single dose inducing likely life-long protection via establishment of non-sterilizing immunity.(2,3) Despite the success of the YF-17D vaccine, production of vaccine stocks has been challenging due to the reliance on cell-based production techniques, limited viral seed stocks due to propagation using an inconsistent process that relies on a limited supply of pathogen-free chicken eggs, and batch-to-batch vaccine production inconsistencies.(4–6) Limited production in recent years and increasing outbreaks have led to a near total depletion of global YF-17D vaccine stockpiles, resulting in dose-reducing measures to stretch the supply.(7,8) Additionally, YF-17D is contraindicated in persons with compromised immune systems, and infrequently can cause disseminated disease in the elderly and young infants.(9)

ZIKV is found in tropical regions of Africa, Asia, and the Americas, with ZIKV infections potentially leading to fever, rash, joint pain, and conjunctivitis.(10) Additionally, ZIKV is one of the few arboviruses to display both horizontal (including sexual contact) and vertical (maternal-fetal) transmission in humans, and can lead to significant birth defects when contracted during pregnancy.(11) Both YFV and ZIKV pose a significant public health threat and have caused outbreaks and epidemics since 2000 in multiple areas of the world.(12) No approved ZIKV vaccine exists despite the development of vaccine candidates that span a range of traditional and next-generation vaccine technologies.(13) Recent Phase 1 clinical trials have demonstrated an excellent safety profile for ZIKV mRNA vaccines, but some of these candidates did not establish a sufficient immune response.(14)

A new vaccine platform would be of great use to fully address the global public health issues caused by these flaviviruses.(15) Ideal next-generation flavivirus vaccines would provide levels of protection against disease comparable to that induced by YF-17D, allow for adaptation to emerging viral strains, be manufacturable at a low cost with a reliable and consistent process, and be sufficiently thermostable for global distribution. Moreover, given the geographical overlap between viruses spread by *Aedes* mosquitoes like YFV, ZIKV, dengue, chikungunya, and others, the ability to create multivalent arbovirus vaccine candidates would be of great benefit.

We previously developed a vaccine platform that delivers self-amplifying RNA (saRNA) to target cells via a simple and unique nanostructured lipid carrier (NLC) delivery system.(16) These saRNA-NLC vaccine formulations are straightforward to manufacture at scale and readily lyophilizable, increasing shelf life to at least 21+ months when refrigerated and up to 6 months at room temperature.(17) This technology has been applied in the development of a clinical-stage SARS-CoV-2 vaccine, among others.(16–21) Here, we created saRNA-NLC vaccine candidates against YFV (AAHI-YFV) and ZIKV (AAHI-ZKV) using this established platform, and, in C57BL/6 mice and Syrian golden hamsters, illustrate the ready adaptability of the vaccine platform to support bivalent immunization. The monovalent and bivalent vaccines induced robust humoral and cellular flavivirus-specific immune responses and conferred complete protection against viral challenge, demonstrating the feasibility of multivalent saRNA-NLC vaccines against mosquito-borne diseases.

## METHODS

### Protein modeling

Protein modeling and analysis of the YF-17D (RCSB PDB ID: 6IW4) and ZIKV envelope (E) (RCSB PDB ID: 5JHM) domains were conducted in PyMOL 2.0 (Schrödinger, LLC). Protein alignments and structural similarity evaluation were done using the Research Collaboratory for Structural Bioinformatics (RCSB) Protein Data Bank (PDB) pairwise alignment server under jFATCAT rigid alignment.(22–24)

### saRNA cloning and production

YFV pre-membrane-envelope (prM-E) (GenBank ID: JN628281.1), ZIKV prM-E (GenBank ID: KJ776791), and secreted alkaline phosphatase protein (SEAP) (GenBank ID: LC380029.1) inserts were subcloned into T7-VEEVRep plasmids.(16,17) Linearized DNA plasmids were used to synthesize YFV prM-E, ZIKV prM-E, and SEAP saRNAs using a previously optimized *in vitro* transcription and purification protocol(16,17) (see Supplementary Methods).

### NLC production

The NLC formulation consisted of an oil core stabilized within an aqueous buffer using appropriate surfactants and was produced by microfluidization as previously described (17) (see Supplementary Methods).

### Vaccine complexing and characterization

We generated saRNA-NLC vaccine complexes by simple mixing (17,19) (see Supplementary Methods). Dynamic light scattering (DLS) using the Zetasizer Nano ZS (Malvern Panalytical) was used for nanoparticle size determination as previously described. (16,17,19) The resulting intensity-weighted Z-average diameter was determined for each formulation and averaged from three measurements per formulation.

### Transfection, protein harvest, and western blotting

HEK293T cells (American Type Culture Collection (ATCC) #CRL-11268) were transfected with complexed vaccine. YFV and ZIKV E protein expression was then measured by western blot of transfected cell lysates (see Supplementary Methods).

### Animal studies

#### Mouse studies

All animal work was done under the oversight of the Bloodworks Northwest Research Institute’s (Seattle, WA) Institutional Animal Care and Use Committee (IACUC), protocol #5389-01. All animal work followed applicable sections of the Final Rules of the Animal Welfare Act regulations (9 CFR Parts 1, 2, and 3) and the *Guide for the Care and Use of Laboratory Animals, Eighth Edition*.

C57BL/6J mice obtained from The Jackson Laboratory or Charles River Laboratories were used for mouse studies. Mice were 6-8 weeks of age at study onset, and groups were sex balanced. Mice were immunized with saRNA-NLC vaccines by intramuscular injection bilaterally in the rear quadriceps muscle (50 µL/leg, 100 µL total) for a prime (Day 0) and a boost (for some groups) 4 weeks later (Day 28). YF-17D was sourced from the University of Texas Medical Branch World Reference Center for Emerging Viruses and Arboviruses and propagated for >3 passages in Vero cells (ATTC #CCL-81). For mice vaccinated with YF-17D, one 20 µL dose containing 10^4^ PFU was administered via subcutaneous rear footpad injection. Serum and spleen samples were collected in-life and at terminal harvest (see Supplementary Methods).

Mice challenged with ZIKV were given 3 mg interferon alpha receptor (IFNAR) blocking monoclonal antibody (MAR1-5A3, BioXCell #BE0241) via intraperitoneal injection on challenge day -1 (D29 or D57), as previously described.(16,25) Additional 1 mg doses were administered on challenge day +1 and +4 (D30 and D33 or D59 and D62). Mice were weighed immediately prior to and daily during the 15-day post-prime and post-boost ZIKV challenge periods (D30-51 and D58-79). Mice were challenged with 10^6^ PFU of ZIKV (strain: MA-Zika Dakar; see Supplementary Methods for virus culture) via bilateral subcutaneous injection (20 µL/rear footpad, 40 µL total).(26) Excess viral stocks used on each challenge were assessed by plaque assay to verify viral titer (actual titer administered = 8.8×10^5^ PFU).

#### Hamster studies

All animal care and husbandry were conducted according to protocols approved by the Utah State University IACUC, protocol #10010 (see Supplementary Methods).

Syrian golden hamsters, weighing 80-90 g (approximately 34-40 days old), were purchased from Charles River Laboratories, and randomly divided into sex-balanced groups. Hamsters were immunized with saRNA-NLC vaccines as described above for mice. For YF-17D vaccinated hamsters, a 1-passage stock of human vaccine YF-VAX (Sanofi) was diluted in sterile saline, and 100 µL was administered via subcutaneous injection into the inguinal fold for each immunization. A cohort of hamsters was also left unvaccinated and uninfected as normal health control animals. Serum samples were collected in-life and at terminal harvest (see Supplementary Methods).

Hamsters were weighed immediately prior to and daily during the 21-day post-prime and post-boost YFV challenge periods (D30-51 and D58-79). Hamsters were challenged with 200 CCID_50_ (50% cell culture infectious dose) of hamster-adapted YFV (strain: Jimenez; see Supplementary Methods for virus culture) via bilateral intraperitoneal injection of 100 µL (200 µL total).(27)

### ELISA

Serum YFV and ZIKV E protein-binding IgG antibodies were measured by ELISA (see Supplementary Methods).(19) Briefly, plates were coated with 1 µg/mL of recombinant YFV E protein (Meridian Bioscience #R01709) or recombinant ZIKV E protein (Meridian Bioscience #R01635) in PBS and incubated overnight at 4°C. A pan-flavivirus E protein-binding monoclonal antibody (4G2; Novus Biologicals #NBP2-52709) was used as the positive control.

### Plaque reduction neutralization test

Serum YFV and ZIKV neutralizing antibody (nAb) titers were determined by a 50% plaque reduction neutralization test (PRNT_50_)(see Supplementary Methods).(17) Briefly, PRNTs were performed with YF-17D or ZIKV FSS13025 viruses.

### Flow cytometry

Intracellular cytokine staining and flow cytometry were conducted to measure antigen-reactive splenic T cells (see Supplementary Methods) as previously described.(19)

### Viral titer measurements

See Supplementary Methods.

### Alanine aminotransferase assay

See Supplementary Methods.

### Scientific rigor

See Supplementary Methods for discussion of replicates, sample size estimations, randomization, blinding, and inclusion and exclusion criteria.

### Statistical analyses

Statistical analysis was conducted using GraphPad Prism 10.3.0. To evaluate immunogenicity data, we examined distribution and variance through Q-Q plot and residual plot analysis. For serum E protein-specific IgG, nAb titers, and viremia, statistics were performed on log_10_ transformed data using one-way ANOVA with Dunnett’s correction or Tukey’s correction for multiple comparisons, or mixed-effect analysis with Tukey’s correction. For weight change, alanine aminotransferase (ALT), and antigen-reactive T cells, statistics were performed using one-way ANOVA with Dunnett’s correction, Brown-Forsythe and Welch ANOVA with Dunnett’s T3 correction, or Kruskal-Wallis with Dunn’s correction for multiple comparisons. For survival curves, statistics were performed by Mantel-Cox log-rank test.

## RESULTS

### Design of saRNA-NLC vaccines against YFV and ZIKV

We developed YFV and ZIKV saRNA-NLC vaccines for delivery in monovalent or bivalent formulations. Each saRNA contains two open reading frames (ORFs), a 3’ polyadenylation sequence (42 As encoded into the DNA template), and a 5’ cap-0 structure (**Figure 1a**). The first ORF encodes the non-structural proteins (nsPs) from the TC-83 strain of Venezuelan equine encephalitis virus (VEEV)(28), and the second ORF encodes codon-optimized prM and E genes from the genomes of YF-17D or ZIKV A/PF/2013, respectively. The E protein of each virus represents the primary viral antigen, while the prM domain stabilizes the E protein for proper folding during translation.(29,30) YFV and ZIKV saRNA constructs were complexed as monovalent formulations with AAHI’s unique NLC delivery particle (**Figure 1b**). The cationic NLC allows for strong electrostatic binding of the saRNA to the outside of the particle, resulting in vaccine particles approximately 80 nm in diameter (**Supplementary Figure 1).** The YFV and ZIKV E domains share significant (40%) protein sequence similarity and 97% structural conservation (**Figure 1c**), suggesting that they should have similar expression behavior and cellular processing, and may elicit comparable immune responses. The formulated vaccines were then transfected into HEK293T cells, in which YFV or ZIKV E protein expression was detected 24 hours post-transfection by western blot (**Figure 1d**). Given the similarities in both biophysical properties of the saRNA-NLC complexes and the antigenic proteins expressed, we expected the YFV and ZIKV vaccine candidates to have similar patterns of immunogenicity.

**Figure 1.**
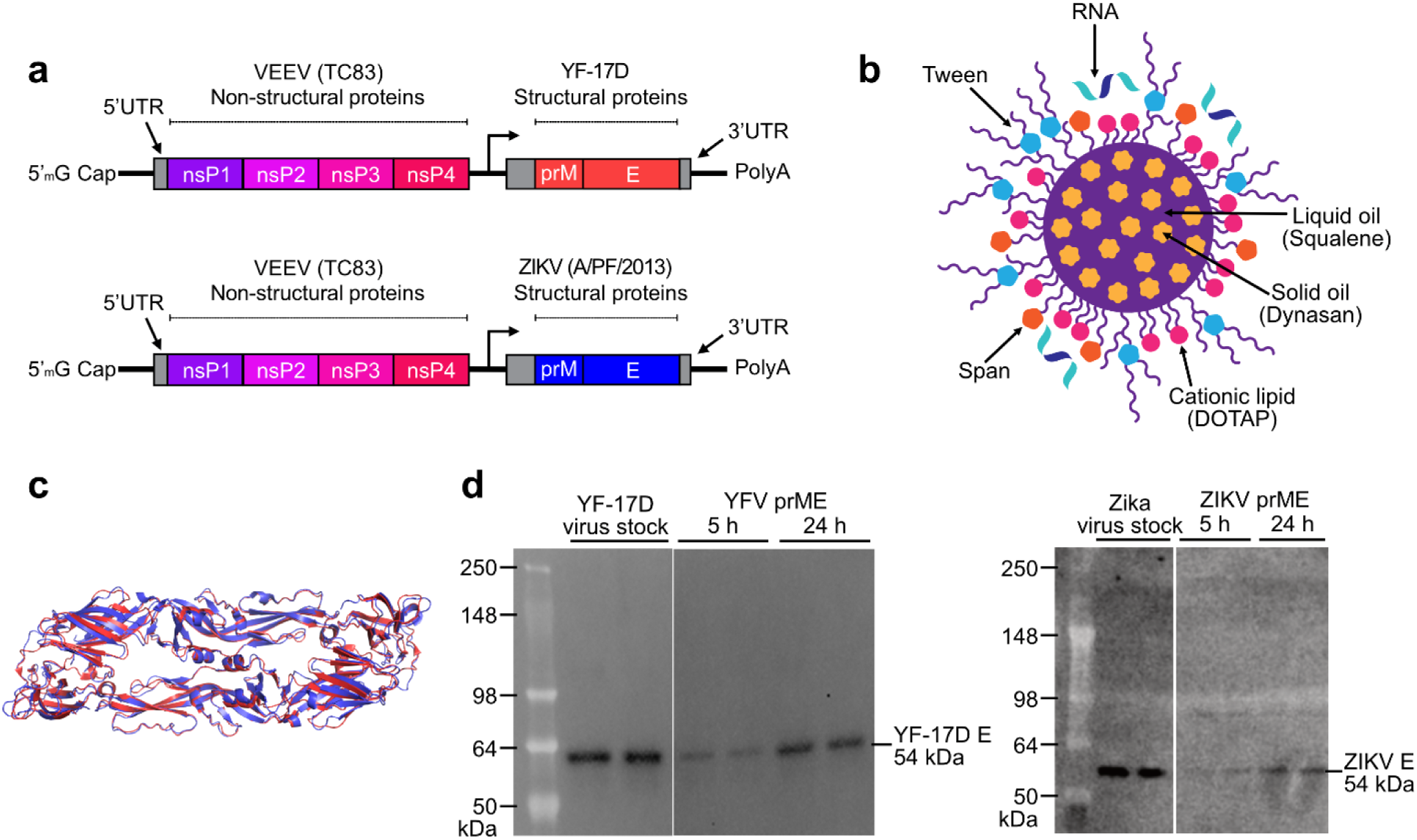
Characterization of the components in AAHI’s bivalent flavivirus saRNA-NLC vaccine. **(a)** Construct designs for the YFV and ZIKV saRNA vaccine constructs. YFV replicon size is 9.89 kb, and ZIKV replicon size is 9.94 kb. **(b)** Schematic of the NLC RNA delivery particle. Design by Cassandra Baden. **(c)** Structural alignment of the YF-17D (red) and ZIKV (blue) E proteins, indicating significant structural similarity between the two antigens. **(d)** Western blot verifying *in vitro* protein expression of 54 kDa YFV and ZIKV E protein in cellular lysates after vaccine HEK293T cell transfection. YF-17D and Zika virus stocks were diluted to 10⁴ PFU/lane and run in duplicate.

### A monovalent YFV saRNA vaccine induces robust serum antibody responses in mice

Serum antibody titers are a primary correlate of protection (CoP) against YFV infection.(2,3) Thus, a viable YFV vaccine candidate should establish robust and long-lasting nAb responses, ideally similar to those induced by the approved YF-17D vaccine.(2) To study the ability of our monovalent YFV saRNA-NLC vaccine (AAHI-YFV) to induce robust nAb responses, C57BL/6 mice were intramuscularly primed (Day 0) and boosted (Day 28) with various doses (5, 10, 20, and 30 µg) of our monovalent YFV vaccine (AAHI-YFV). Mice immunized with a single dose (10^4^ PFU) of YF-17D were used as a positive vaccine control group. Mice injected with 10 µg of saRNA-NLC expressing the non-immunogenic protein SEAP were used to control for induction of non-specific immune responses.

Serum YFV E protein-specific antibody titers in AAHI-YFV vaccinated mice showed little to no dose-dependency above 10 µg saRNA (**Figure 2a-b**). While a 5 µg saRNA dose induced more heterogeneous serum IgG titers, high serum IgG titers were observed in mice immunized with 10 to 30 µg of saRNA, equivalent to those induced by vaccination with 10^4^ PFU of YF-17D.

**Figure 2.**
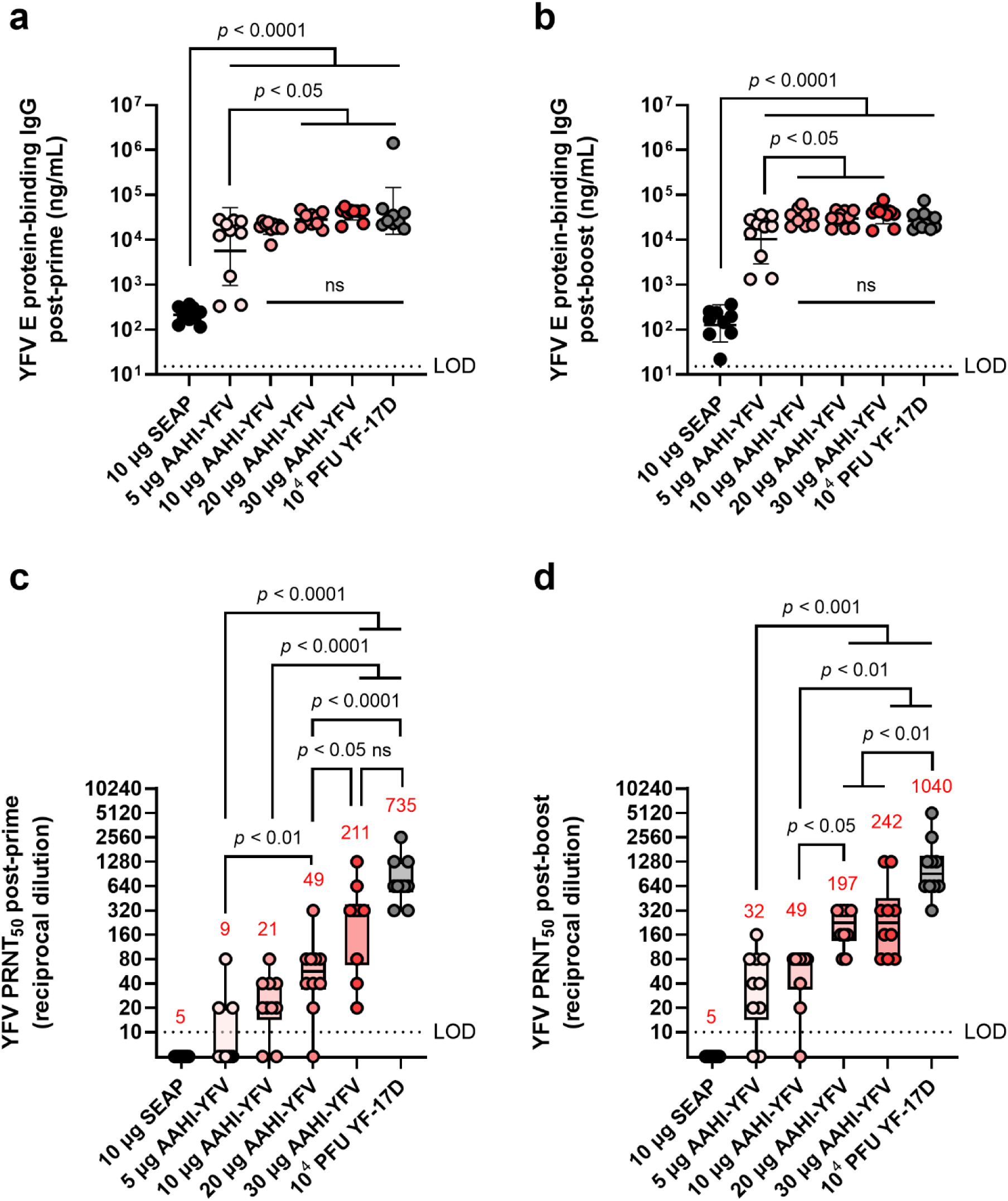
YFV saRNA-NLC vaccination induces serum antigen-binding IgG and neutralizing antibody responses in mice comparable to those induced by the YF-17D vaccine. Mice were vaccinated with a dose range of AAHI-YFV saRNA-NLC and **(a-b)** serum YFV E protein-binding IgG and **(c-d)** YFV neutralizing antibody titers were measured **(a, c)** 28 days post-prime and **(b, d)** 28 days post-boost. The YF-17D group represents mice that were subcutaneously dosed once with 10⁴ PFU of YF-17D. Statistical analysis was conducted on log_10_ transformed data using one-way ANOVA with Tukey’s correction for multiple comparisons. ns = non-significant (*p* > 0.05). Black dotted line shows the limit of detection (LOD) for the assay. Results are from a single independent experiment; *n* = 10 mice per group, 5 male and 5 female. **(a-b)** Scatter plots show geometric mean ± geometric SD. **(c-d)** Box plots show median and IQR ± min/max value. Red numerical values represent group PRNT_50_ GMT.

We observed a strong dose-dependency in serum YFV nAb titers (**Figure 2c-d**). All AAHI-YFV dosing groups, apart from the 5 µg group, generated a PRNT_50_ geometric mean titer (GMT) ≥20 post-prime and ≥40 post-boost, which exceeded the CoP established in other models: PRNT_50_ ≥10 (humans), PRNT_50_ ≥40 (hamsters), and PRNT_50_ ≥20 (non-human primates).(2,3,31,32) Post-boost, both 5 and 10 µg vaccinated groups generated a PRNT_50_ GMT ≥20, though some individual mouse PRNT_50_ titers remained below the limit of detection. Conversely, all mice vaccinated with 20 and 30 µg doses developed PRNT_50_ titers ≥80 post-boost. These high-dose vaccinated groups had a post-boost nAb GMT approaching but not equaling those induced by YF-17D vaccination. The post-boost PRNT_50_ titers established by all saRNA doses exceeded the CoP for other models, consistent with the post-prime results, and reached levels indicative of protection in other mouse model studies(2,33,34). Overall, these data demonstrate that AAHI-YFV induces potent humoral immunity in mice, particularly at 10 to 30 µg doses.

### YFV saRNA vaccination establishes durable humoral immune responses at higher doses

The FDA-approved YF-17D vaccine provides life-long protection against disease for most individuals.(35) We previously demonstrated that our ZIKV saRNA-NLC vaccine generates a durable serum nAb response up to 7 months post-prime and post-boost immunization.(16) Therefore, we investigated the longevity of humoral immunity following AAHI-YFV vaccination. To do this, prime-boost 10 µg, 20 µg, and 30 µg AAHI-YFV and YF-17D vaccinated C57BL/6 mice from the AAHI-YFV dosing study above were maintained for 6 months post-boost to investigate antibody durability.

As expected, serum antigen-binding IgG titers declined significantly over the 6 months post-boost vaccination but remained well in the detectable range for all vaccinated groups (**Figure 3a**). Notably, mice immunized with all doses of the AAHI-YFV saRNA-NLC vaccine showed serum IgG titers at 6 months post-boost comparable to YF-17D-vaccinated mice (10 µg: *p* = 0.99, 20 µg: *p =* 0.99, and 30 µg: *p =* 0.58) suggesting that the saRNA-NLC vaccine can indeed induce durable humoral immune responses to YFV. Serum nAb titers showed similar results to antigen-binding IgG titers, waning over the 6-month period post-boost but maintaining largely detectable nAb titers in the saRNA-NLC vaccinated mice (**Figure 3b**). Furthermore, at 6 months post-boost, mice receiving 20 and 30 µg of AAHI-YFV maintained a nAb GMT commensurate to that induced by YF-17D (20 µg: *p* = 0.1342 and 30 µg: *p* = 0.9540). Interestingly, while the nAb GMT of the YF-17D group was greater than the 30 µg AAHI-YFV group after the initial vaccinations (**Figure 2d**), the nAb GMT of the YF-17D group fell more precipitously and was equivalent to the 30 µg AAHI-YFV group at 6 months post-boost (**Figure 3b**). Critically, both groups maintained nAb titers well above the neutralizing CoP for humans (PRNT_50_ ≥10) at the final timepoint.(3,36) Together, these data indicate that the potent antibody response induced in mice by our AAHI-YFV vaccine is durable, as serum nAb titers were maintained at levels similar to YF-17D vaccination 6 months after the original vaccination series.

**Figure 3.**
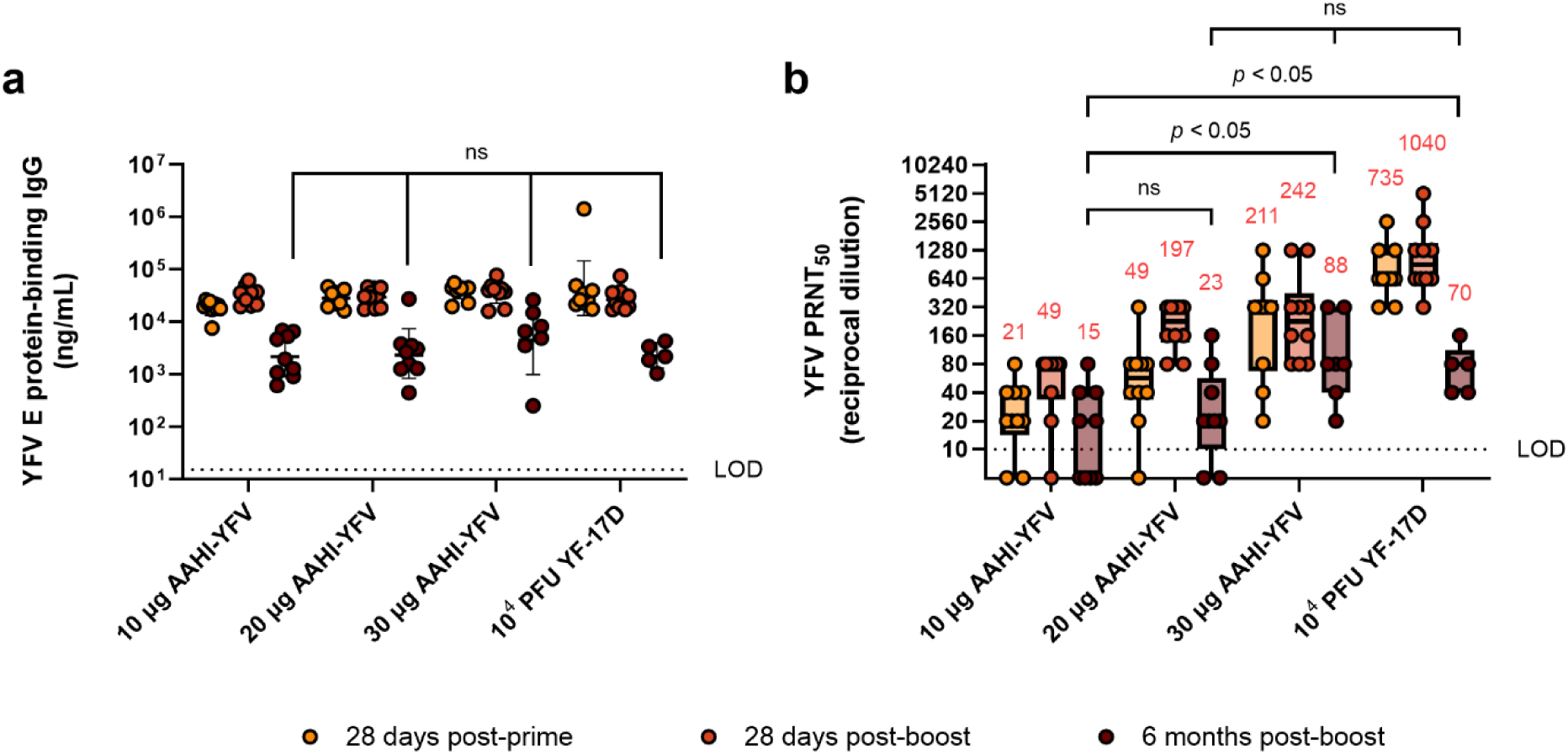
Durability of YFV-specific responses from monovalent saRNA-NLC vaccination. **(a)** Serum YFV E protein-binding IgG and **(b)** YFV neutralizing antibody responses were assessed 28 days post-prime, 28 days post-boost, and 6 months post-boost with AAHI-YFV saRNA-NLC vaccines. The YF-17D group represents mice that were subcutaneously dosed once with 10⁴ PFU of YF-17D. Statistical analysis was conducted on log_10_ transformed data using a mixed-effect analysis with Tukey’s correction for multiple comparisons. Black dotted line shows the limit of detection (LOD) of the assay. ns = non-significant (*p* > 0.05). Results are from a single independent experiment; *n* = 10 mice per group, 5 male and 5 female. Animals were removed from the 6-month time point due to length of study: 1 animal from the 10 µg group, 1 animal from the 20 µg group, 3 animals from the 30 µg group, and 5 animals from the YF-17D group. **(a)** Scatter plots show geometric mean ± geometric SD. **(b)** Box plots show median and IQR ± min/max value. Red numerical values represent group PRNT_50_ GMT.

### A ZIKV saRNA monovalent vaccine establishes a potent nAb response at various doses

Like other flaviviruses, nAbs to ZIKV are thought to be the best CoP from infection.(37) We therefore evaluated the antibody response to our monovalent ZIKV vaccine (AAHI-ZKV) as we did for AAHI-YFV. C57BL/6 mice were prime or prime-boost immunized with escalating doses (0.1, 1, 10, and 30 µg) of AAHI-ZKV, representing a lower focused dose range than tested for the YFV vaccine candidate due to the previously determined high potency of the ZIKV saRNA-NLC vaccine.(16)

We observed a dose-dependent increase in ZIKV E protein-specific serum IgG. A dose of 0.1 µg AAHI-ZKV generated responses comparable to background seen in SEAP-expressing vector control saRNA-NLC dosed mice, whereas mice primed and boosted with 10 or 30 µg of saRNA developed the highest serum IgG titers (**Figure 4a-b**), as was also observed for AAHI-YFV.

**Figure 4.**
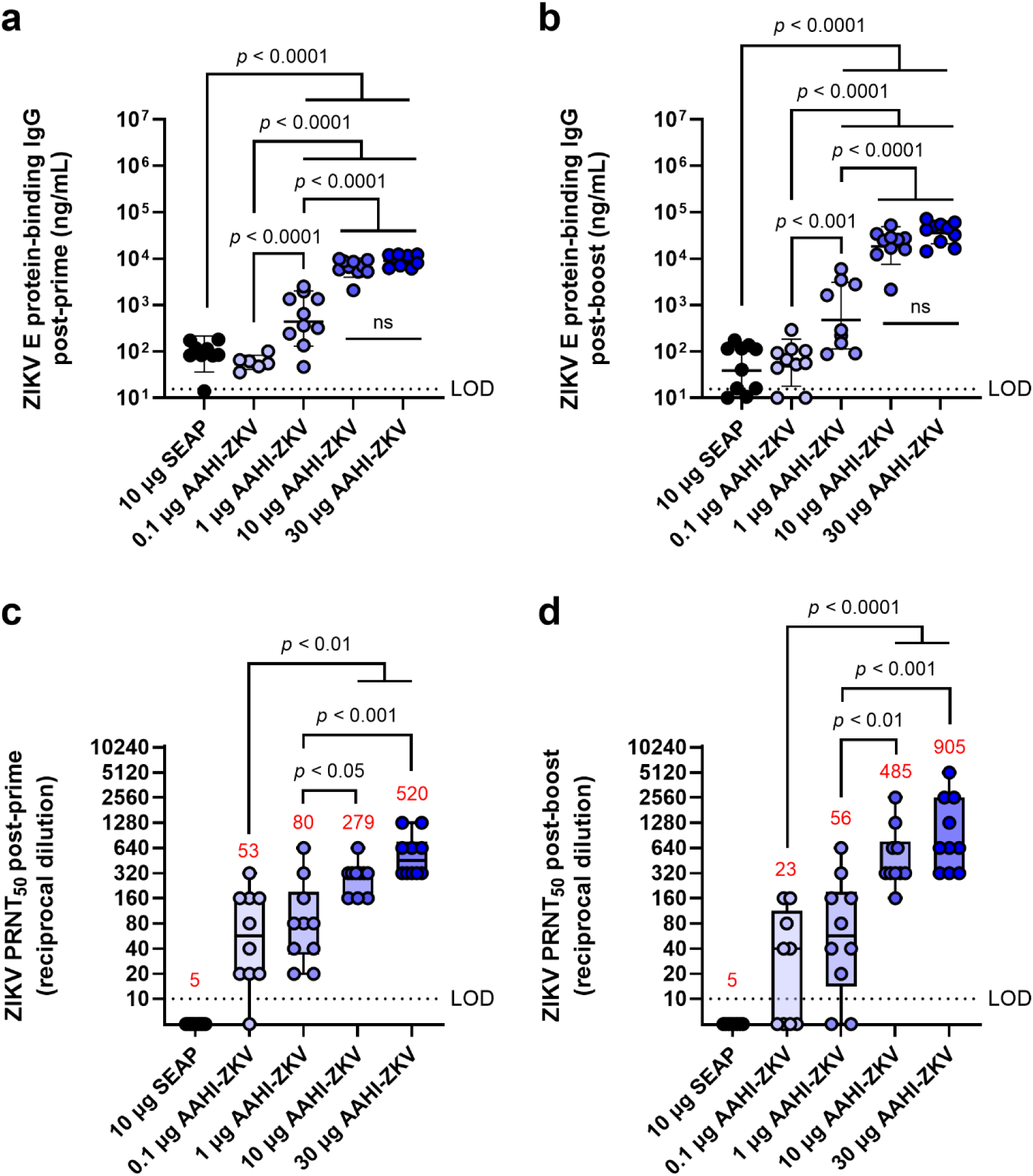
ZIKV-specific serum IgG and neutralizing antibody responses following monovalent saRNA-NLC vaccination. Mice were vaccinated with a dose range of AAHI-ZKV saRNA-NLC and **(a-b)** serum ZIKV E protein-binding IgG and **(c-d)** ZIKV neutralizing antibody titers were measured **(a, c)** 28 days post-prime and **(b, d)** 28 days post-boost. Statistical analysis was conducted on log_10_ transformed data using one-way ANOVA with Tukey’s correction for multiple comparisons. Black dotted line shows the limit of detection (LOD) for the assay. ns = non-significant (*p* > 0.05). Results are from a single independent experiment; *n* = 10 mice per group, 5 male and 5 female. **(a-b)** Scatter plots show geometric mean ± geometric SD. **(c-d)** Box plots show median and IQR ± min/max value. Red numerical values represent group PRNT_50_ GMT.

We also noted a dose-dependent increase in serum nAb titers. Despite background levels of ZIKV antigen-binding IgG at lower doses of saRNA, 0.1 and 1 µg vaccine doses induced a ZIKV PRNT_50_ GMT ≥10 (**Figure 4c-d**). This nAb titer was previously found to confer complete protection against lethal ZIKV challenge in mice,(16) despite being lower than the CoP identified through passive transfer studies in mice (PRNT_50_ GMT ≥100).(38,39) Whereas low-dose AAHI-ZKV vaccination induced variable PRNT_50_ titers in individual animals, 10 and 30 µg doses induced PRNT_50_ titers ≥160 in all mice both post-prime and post-boost. Together, these data demonstrate that both the AAHI-YFV and AAHI-ZKV are highly immunogenic, particularly at the higher saRNA doses tested, with no deleterious outcomes.

### Bivalent vaccine formulations induce nAbs to both targets comparable to monovalent formulations

After verification of the monovalent formulations, we next evaluated mixing strategies for bivalent saRNA-NLC vaccine administration using 10 µg doses of each saRNA. Three bivalent vaccine dosing strategies were tested, to determine optimal bivalent vaccine manufacture processes and to evaluate the presence of any immune interference between the two flaviviral antigen-expressing saRNA species. These mixing strategies included (1) delivering individual monovalent saRNA vaccines in anatomically distinct locations represented by separate legs (“Split monovalent”), (2) mixing pre-formed monovalent YFV and ZIKV saRNA-NLC vaccines (“Complex -> Mix”) with each vaccine complex delivering a single saRNA species, or (3) mixing both vaccine saRNAs prior to complexing with NLCs (“Mix -> Complex”), resulting in vaccine complexes delivering both saRNA species (**Supplementary Table 1**).

Alone, monovalent vaccines induced serum nAb responses against their respective flavivirus as expected, with little to no evidence of cross-neutralization (**Figure 5)**. All mice receiving any of the bivalent vaccines (AAHI-YFV/ZKV), regardless of how the bivalent vaccine was created and injected, induced statistically equivalent nAb titers to those induced by the monovalent formulations. No evidence of any immune interference between the two saRNA-expressed flavivirus antigens was noted, regardless of the bivalent vaccine mixing strategy.

**Figure 5.**
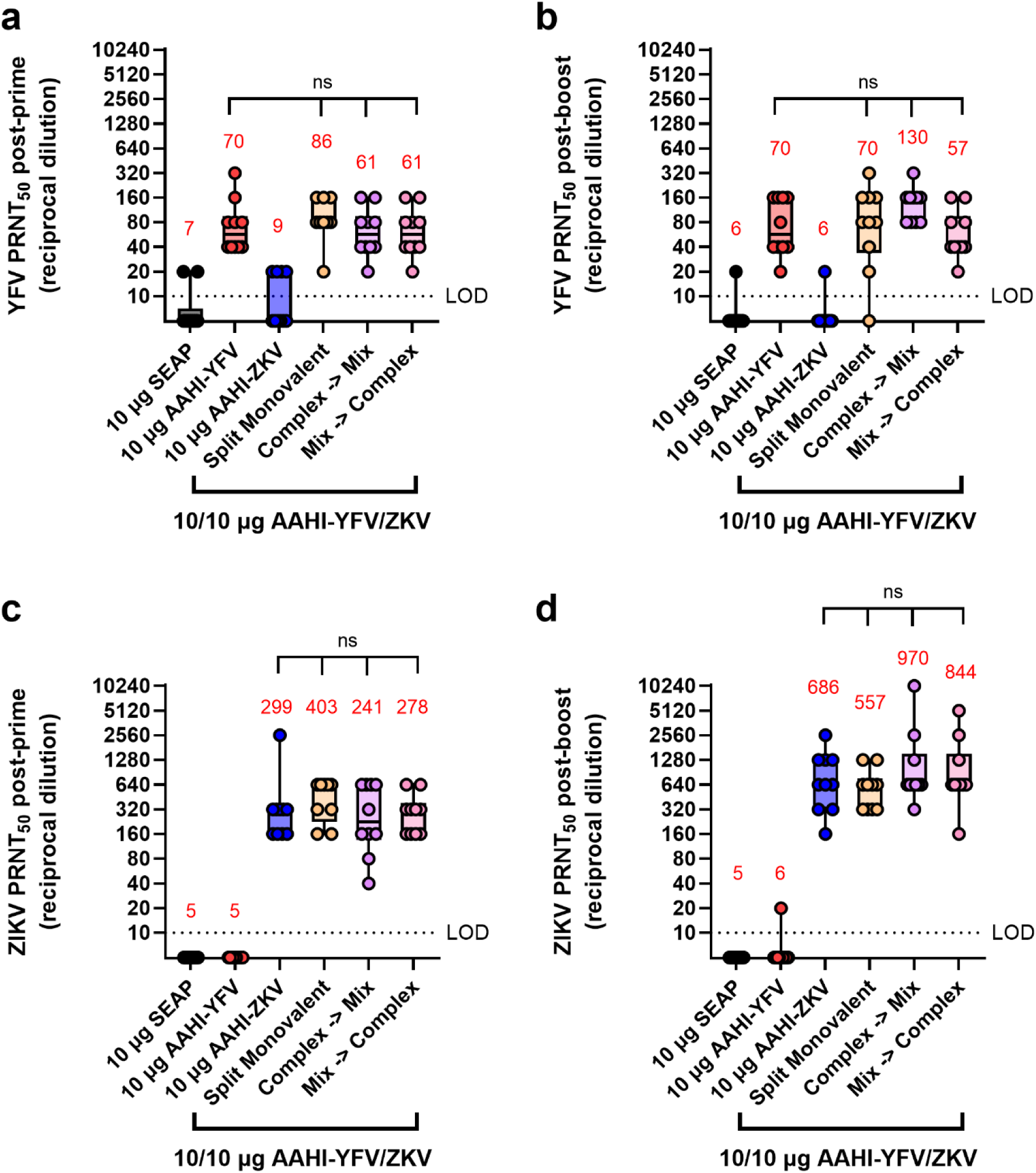
Serum YFV and ZIKV neutralizing antibody titers resulting from different bivalent vaccination strategies. Mice were vaccinated with monovalent or bivalent formulations of AAHI-YFV or AAHI-ZKV saRNA-NLC and serum neutralizing antibody titers against **(a, b)** YFV and **(c, d)** ZIKV were measured **(a, c)** 28 days post-prime and **(b, d)** 28 days post-boost. Statistical analysis was conducted on log_10_ transformed data using one-way ANOVA with Tukey’s correction for multiple comparisons. Comparisons were made against monovalent AAHI-YFV or AAHI-ZKV and between bivalent AAHI-YFV/ZKV groups. ns = non-significant (*p* > 0.05). Black dotted line shows the limit of detection (LOD) for the assay. Results are from a single independent experiment; *n* = 10 mice per group, 5 male and 5 female. Box plots show median and IQR ± min/max value. Red numerical values represent group PRNT_50_ GMT.

### Bivalent vaccine dosing establishes robust antigen-responsive CD4^+^ and CD8^+^ T cell populations

We next studied the cellular responses to the different bivalent dosing strategies investigated in the above mouse study, looking at the antigen-specific responses of splenic CD4^+^ and CD8^+^ T cells. In mice, individual antigen-specific T cells are estimated to vary in frequency from 20 – 200 cells per ∼2 x 10^7^ naïve CD4^+^ or CD8^+^ T cells (0.0001 – 0.001%).(40,41) Levels of splenic YFV and ZIKV antigen-reactive activated (CD44^+^) polyfunctional CD4^+^ T cells, which are those simultaneously expressing IFN-γ^+^, IL-2^+^, and TNF-α^+^ in response to antigen stimulation, were assessed by flow cytometry. We also examined the antigen-reactive IFN-γ^+^ CD8^+^ CD44^+^ T cell responses in these vaccinated mice.

Flavivirus E protein-reactive polyfunctional CD4^+^ T cells were readily detectable (>0.5% of total CD4^+^ CD44^+^ T cells) post-prime and post-boost in AAHI-YFV or AAHI-ZKV vaccinated mice. Polyfunctional CD4^+^ T cells were not detected in mice vaccinated with the SEAP-expressing saRNA-NLC negative control (**Figure 6a-d**). Like the PRNT_50_ data, splenic CD4^+^ T cells in groups immunized with monovalent vaccines were only reactive to the corresponding antigen peptide pools, indicating no induction of cross-reactive CD4^+^ T cells by the two flavivirus vaccines. All bivalent doses, regardless of mixing and dosing strategies, elicited comparable levels of YFV- and ZIKV-reactive polyfunctional CD4^+^ T cells with respect to each other and corresponding monovalent groups, both post-prime and post-boost.

**Figure 6.**
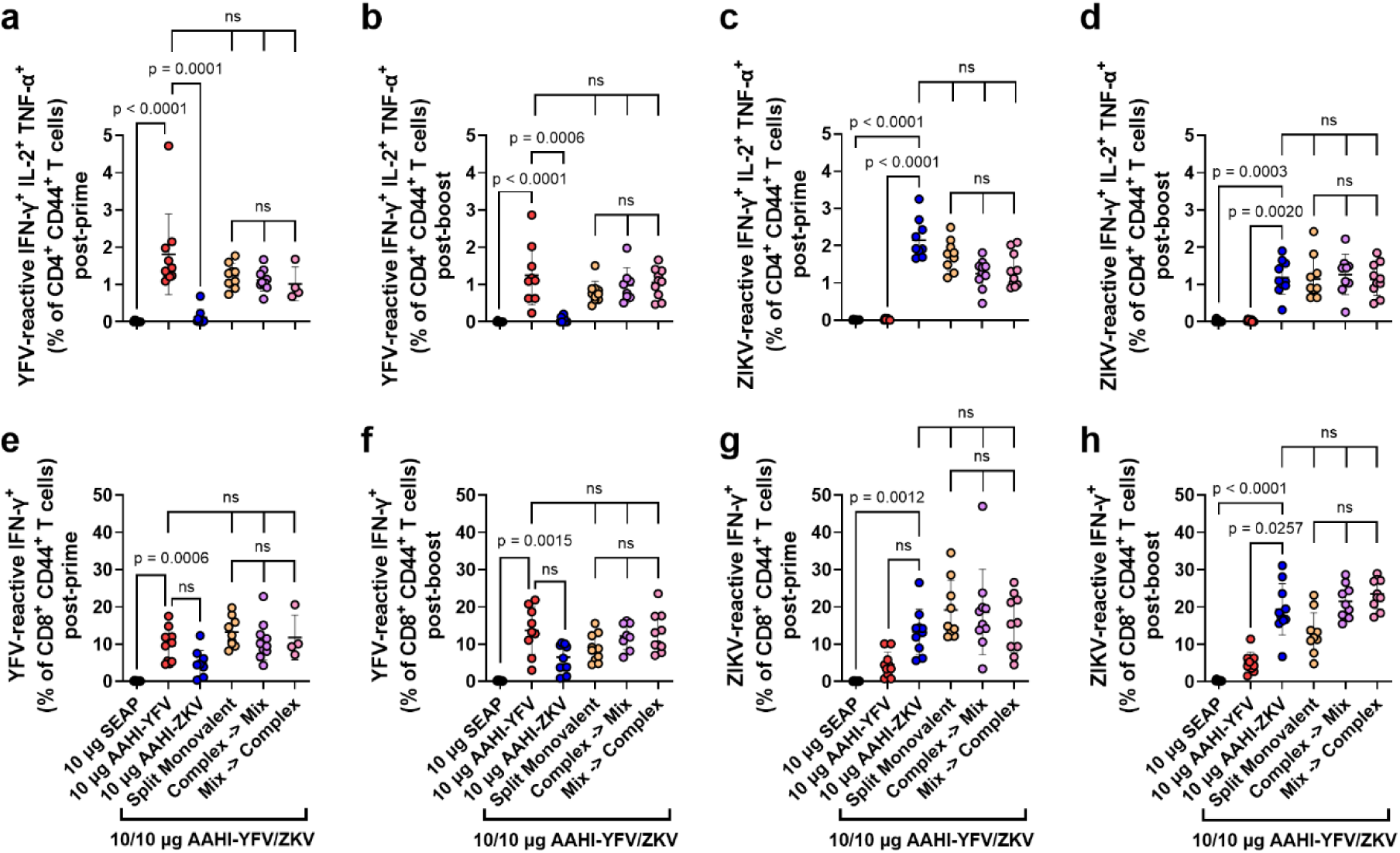
Monovalent and bivalent YFV and ZIKV saRNA-NLC vaccination induces polyfunctional CD4^+^ and IFN-γ^+^ CD8^+^ T cells. Mice were vaccinated with monovalent or bivalent formulations of AAHI-YFV or AAHI-ZKV saRNA-NLC, and splenocytes were isolated **(a, c, e, g)** 28 days post-prime or **(b, d, f, h)** 28 days post-boost. The antigen-specific polyfunctional (IFN-γ^+^ IL-2^+^ TNFα^+^) CD4^+^ T cell response was measured following stimulation with (**a, b**) YFV or (**c, d**) ZIKV prM-E peptide pools. The antigen-specific IFN-γ^+^ CD8^+^ T cell response was measured following stimulation with (**e, f**) YFV or (**g, h**) ZIKV prM-E peptide pools. Statistics were measured using Kruskal-Wallis with Dunn’s correction (**a-e, g, h**) or Brown-Forsythe and Welch ANOVA with Dunnett’s T3 correction (**f**) for multiple comparisons. Comparisons were made against monovalent AAHI-YFV or AAHI-ZKV and between bivalent AAHI-YFV/ZKV groups. ns= non-significant (*p* > 0.05). Results are from a single independent experiment; *n* = 10 mice per group, 5 male and 5 female. Scatter plots show mean ± SD.

Antigen-reactive IFN-γ^+^ CD8^+^ T cells were readily detectable (>1% of total CD8^+^ CD44^+^ T cells) in all monovalent AAHI-YFV and AAHI-ZKV vaccinated mice post-prime and post-boost (**Figure 6e-h**). In contrast with polyfunctional CD4^+^ T cells, cross-reactive IFN-γ^+^ CD8^+^ T cells were detectable at a percentage approximately half or less than that of the matched vaccine antigen. This observation is not surprising as these two related flaviviruses’ prM-E proteins share significant structural and sequence homology, which appear to include some CD8^+^ T cell epitopes but not CD4^+^ T cell epitopes. All bivalent AAHI-YFV/ZKV dosing strategies elicited comparable levels of IFN-γ^+^ CD8^+^ T cells to those elicited by monovalent vaccines both post-prime and post-boost. Lastly, we saw no consistent differences in YFV-or ZIKV-responsive CD8^+^ T cells between bivalent AAHI-YFV/ZKV groups regardless of the mixing and dosing strategy. Overall, saRNA-NLC immunization leads to robust multi-fold expansion of antigen-reactive T cells, and the mode of formulation and delivery of bivalent vaccines using this platform appears to have minimal effects on either humoral or cellular vaccine immunogenicity in mice, allowing for flexibility in future bivalent saRNA-NLC vaccine formulation strategies.

### Immunization with ZKV saRNA-NLC is protective against lethal ZIKV infection in mice

To evaluate the efficacy of AAHI-ZKV in monovalent and bivalent format, we employed a transiently-immunocompromised mouse ZIKV challenge model.(16,26) C57BL/6 mice were first prime or prime-boost vaccinated with 10 µg monovalent AAHI-YFV or AAHI-ZKV, or 10/10 µg, 5/5 µg, or 1/1 µg bivalent AAHI-YFV/ZKV. A cohort of each study group was evaluated for post-vaccination flavivirus nAb titers and antigen-reactive cellular responses. Thirty days after final vaccination, mice were transiently immunocompromised by injection with α-IFNAR antibody -1, +1, and +4 days post-infection (dpi), and challenged with 10^5^ PFU of MA-Zika Dakar.(16,25) Survival (mortality) and body weight change (morbidity) were assessed as metrics of vaccine efficacy (**Figure 7a and 7e**).

**Figure 7.**
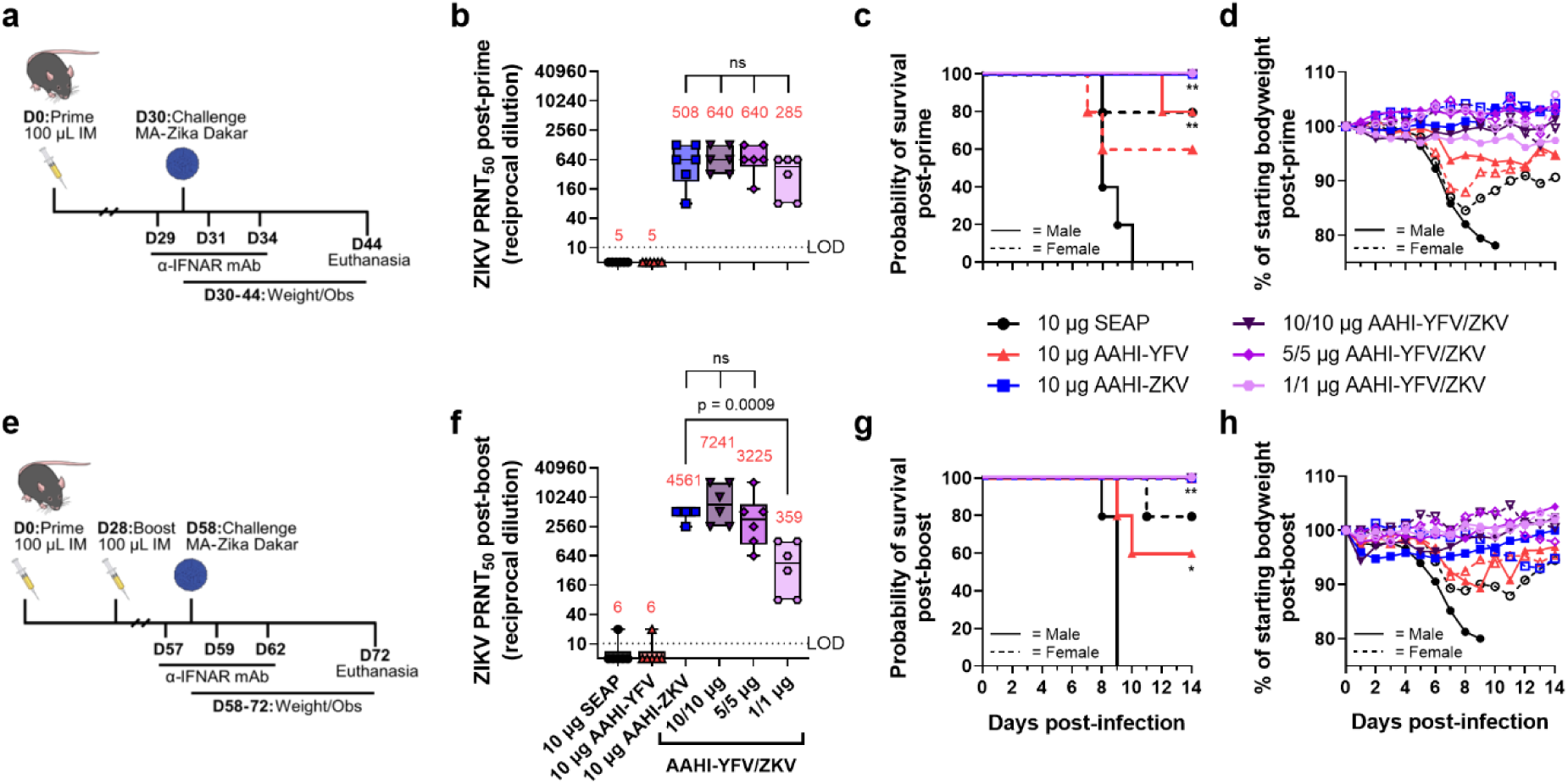
Monovalent and bivalent ZIKV saRNA-NLC vaccination protects mice from lethal ZIKV challenge. (**a**) Post-prime mouse ZIKV challenge study design (81–83) (**b**) Serum ZIKV neutralizing antibody titer 28 days post-prime. (**c**) Post-prime survival curves. (**d**) Post-prime body weight. (**e**) Post-boost mouse ZIKV challenge study design (81–83) (**f**) Serum ZIKV neutralizing antibody titer 28 days post-boost. (**g**) Post-boost survival curves. (**h**) Post-boost body weight. (**b** and **d**) Statistical analysis was conducted on log_10_ transformed data using one-way ANOVA with Dunnett’s correction for multiple comparisons. ns = non-significant (*p* > 0.05). Black dotted line shows the limit of detection (LOD) for the assay. Results are from a single independent experiment; *n* = 6 mice per group, 3 male and 3 female. Box plots show median and IQR ± min/max value. Red numerical values represent group PRNT_50_ GMT. (**c** and **g**) Statistics were assessed by Mantel-Cox log-rank test against 10 µg SEAP-expressing vector control. \**p* < 0.05, \*\**p* < 0.01. Results are from a single independent experiment; *n* = 10 mice per vaccination group, 5 male and 5 female.

Consistent with earlier observations (**Figure 3 and 5**), a single prime dose of AAHI-ZKV, in both monovalent and bivalent formulation, induced robust ZIKV nAb titers well above the predicted CoP identified through passive transfer studies in mice (PRNT_50_ GMT ≥100) (**Figure 7b**) (38,39). A boost dose augmented nAb GMT more than 4-fold in 10 µg monovalent AAHI-ZKV and 10/10 µg and 5/5 µg bivalent AAHI-YFV/ZKV dosed groups (**Figure 7f**). YFV nAb titers were likewise induced by 10 µg AAHI-YFV, and 10/10 µg and 5/5 µg AAHI-YFV/ZKV vaccine doses (**Supplementary Figure 2a**), with limited improvement with a boost dose (**Supplementary Figure 2b**). Lastly, indicative of a systemic vaccine-induced cellular response, splenic antigen-reactive polyfunctional CD4^+^ T cells (>0.2% of total CD4^+^ CD44^+^) (**Supplementary Figure 3a-d**) and CD8^+^ T cells (>0.2% of total CD8^+^ CD44^+^) (**Supplementary Figure 3e-h**) were generated at similar frequencies by both monovalent and bivalent AAHI-ZKV and AAHI-YFV immunization

All mice vaccinated with AAHI-ZKV were completely protected against ZIKV-induced mortality, regardless of valency, dose, or dosing regimen (**Figure 7c and 7g and Supplementary Table 2**). We observed a strong sex-dependent difference in mortality regardless of dosing regimen, with 0% survival in SEAP-expressing vector control vaccinated males but 80% survival in SEAP-expressing vector control vaccinated females. Changes in weight following challenge mirrored survival (**Figure 7d and 7h and Supplemental Figure 4a and 4b**), with sharp decreases in weight noted in SEAP-expressing vector control vaccinated animals, but maintenance of normal body weight 3-6 dpi in AAHI-ZKV immunized mice regardless of sex or dosing regimen. Notably, mice immunized with 10 µg AAHI-YFV were partially protected from ZIKV challenge, showing 60% - 100% survival depending on both sex and dose regimen, and limited weight loss with survivors regaining normal weight by the end of the study.

Together, these results demonstrate (1) strong protection against ZIKV-induced morbidity and mortality by all AAHI-ZKV doses tested regardless of dosing regimen, (2) no detriment of combining AAHI-YFV and AAHI-ZKV into a bivalent vaccine, and (3) significant cross-protection from ZIKV-induced morbidity and mortality conferred by AAHI-YFV immunization.

### Immunization with YFV saRNA-NLC is protective against lethal YFV infection in hamsters

To evaluate the efficacy of AAHI-YFV in monovalent and bivalent format, we employed a hamster YFV challenge model.(42) Syrian golden hamsters were first prime or prime-boost vaccinated with 10 µg monovalent AAHI-YFV or AAHI-ZKV, or 10/10 µg, 5/5 µg, or 1/1 µg bivalent AAHI-YFV/ZKV vaccines. Vaccinated hamsters were evaluated for post-vaccination flavivirus serum nAb titers. Thirty days after final vaccination, hamsters were challenged with 200 CCID_50_ of YFV-Jiménez. Survival (mortality), body weight change and serum ALT (morbidity), and viremia were assessed as metrics of vaccine efficacy (**Figure 8a and 8e**).

**Figure 8.**
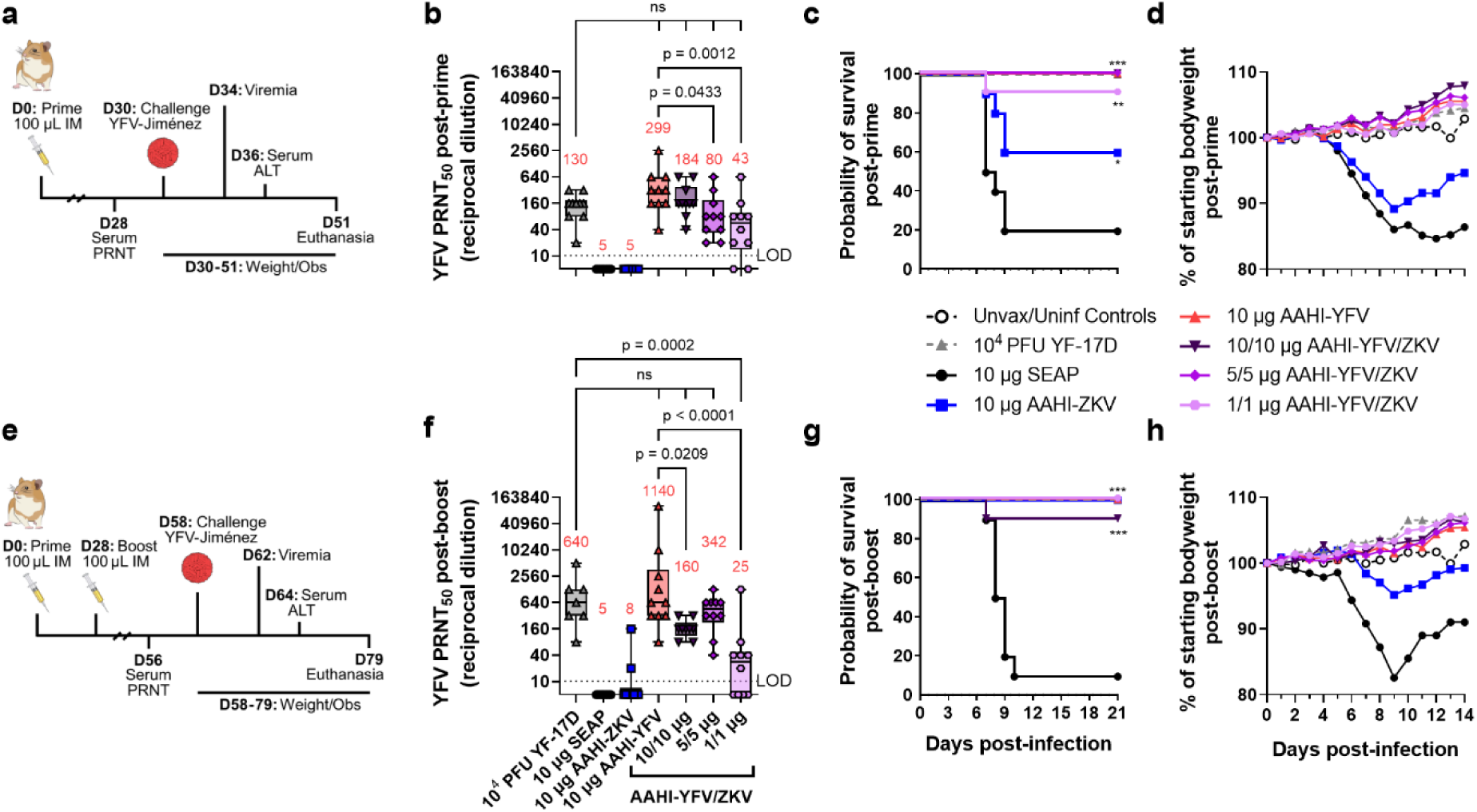
Monovalent and bivalent YFV saRNA-NLC vaccination protects hamsters from lethal YFV challenge. (**a**) Post-prime hamster YFV challenge study design (81,83,84) (**b**) Pre-challenge serum YFV neutralizing antibody titer 28 days post-prime. (**c**) Post-prime survival curves. (**d**) Post-prime body weight. (**e**) Post-boost hamster YFV challenge study design (81,83,84) (**f**) Pre-challenge serum YFV neutralizing antibody titer 28 days post-boost. (**g**) Post-boost survival curves. (**h**) Post-boost body weight. (**b** and **d**) Statistical analysis was conducted on log_10_ transformed data using one-way ANOVA with Dunnett’s correction for multiple comparisons. ns = non-significant (*p* > 0.05). Black dotted line shows the limit of detection (LOD) for the assay. Box plots show median and IQR ± min/max value. Red numerical values represent group PRNT_50_ GMT. (**c** and **g**) Statistics were assessed by Mantel-Cox log-rank test against 10 µg SEAP-expressing vector control. \**p* < 0.05, \*\**p* < 0.01, \*\*\**p* < 0.001. Results are from a single independent experiment; *n* = 10 hamsters per group, 5 male and 5 female, or *n* = 5 female hamsters for unvaccinated/uninfected (Unvax/Uninf) controls.

Consistent with studies in mice, hamsters prime immunized with AAHI-YFV or AAHI-ZKV in both monovalent and bivalent formulation developed group serum nAb GMTs above the predicted CoP for each flavivirus (YFV: PRNT_50_ ≥ 40 in hamsters, ZIKV: PRNT_50_ ≥ 100 in mouse passive transfer studies) (**Figure 8b and Supplementary Figure 5a**).(31,38,39,43) YFV and ZIKV nAb GMTs were generally higher with a prime-boost immunization schedule, though bivalent boosting showed inconsistent improvement for YFV nAb titers in contrast with ZIKV nAb titers (**Figure 8f and Supplementary Figure 5b**). After prime immunization, groups that received a 10/10 µg dose of AAHI-YFV/ZKV had YFV nAb titers comparable to those vaccinated with 10 µg AAHI-YFV. Lastly, serum YFV nAb titers in 10 µg AAHI-YFV and 10/10 µg and 5/5 µg AAHI-YFV/ZKV vaccinated hamsters were comparable to serum YFV nAb titers in YF-17D vaccinated hamsters.

As expected, a SEAP-expressing vector control immunization did not confer protection (20% survival), and YF-17D vaccination was completely protective (100% survival). Protection against mortality after YFV challenge was demonstrated in all study groups that received a prime-only or prime-boost regimen of AAHI-YFV or AAHI-YFV/ZIKV vaccine (**Figure 8c and 8g and Supplementary Table 3**). Prime-only vaccination with AAHI-YFV was highly protective with 100% survival in all doses other than the lowest (1/1 µg) AAHI-YFV/ZKV, which provided significant protection from mortality in 9/10 animals (**Figure 8c**). Interestingly, partial cross-protection was observed in hamsters dosed with 10 µg AAHI-ZKV, resulting in 60% survival (2/5 females and 4/5 males) after prime vaccination that was significantly improved over the SEAP-expressing vector control, and further improved to 100% survival after boost

Morbidity and viremia measurements also revealed a robust protective effect of AAHI-YFV vaccination. Weight loss 3-6 dpi was significantly reduced by prime immunization with AAHI-YFV or YF-17D compared to the SEAP-expressing vector control, whereas AAHI-ZKV vaccination alone did not significantly reduce weight change (**Figure 8d and Supplementary Figure 6a**). Post-boost, however, all AAHI-YFV and AAHI-ZKV vaccination strategies significantly limited weight change 3-6 dpi (**Figure 8h and Supplementary Figure 6d**). Average serum ALT levels 6 dpi, reflecting onset of YFV-induced viscerotropic disease, were at or near baseline in post-prime and post-boost YF-17D, AAHI-YFV, AAHI-ZKV, and AAHI-YFV/ZKV vaccinated hamsters (**Supplementary Figure 6b and 6e**), and were significantly improved relative to SEAP-expressing vector control-vaccinated hamsters. Lastly, viremia 4 dpi was prevented in animals receiving YF-17D, and any monovalent or bivalent dose of AAHI-YFV via prime-only or prime-boost vaccination, with the sole exception of 3/10 animals in the low 1/1 µg AAHI-YFV/ZKV prime-only vaccine dose group (**Supplementary Figure 6c**). Prime-only immunization with 10 µg AAHI-ZKV was insufficient to reduce serum viral titers relative to the SEAP-expressing vector control, but prime-boost AAHI-ZKV immunized animals resulted in reduced viremia relative to controls (**Supplementary Figure 6e**).

In sum, these data demonstrate (1) strong protection against YFV-induced morbidity and mortality provided by AAHI-YFV at all doses tested regardless of dosing regimen, (2) no detriment from bivalent combination of AAHI-YFV and AAHI-ZKV vaccines, and (3) significant cross-protection against YFV infection provided by the AAHI-ZKV vaccine.

## DISCUSSION

The aim of this study was to demonstrate platform feasibility and evaluate the immunogenicity and efficacy of a bivalent saRNA-NLC vaccine, in this case designed to target the geographically overlapping mosquito-borne flaviviruses YFV and ZIKV. Both monovalent AAHI-YFV and AAHI-ZKV immunization induced strong and durable serum flavivirus E protein-binding IgG and virus-specific nAb responses. Similarly robust nAb titers were elicited following bivalent AAHI-YFV/ZKV vaccination, through various dosing strategies and concentrations, at levels equivalent to those induced by the monovalent vaccines. These results were mirrored in the cellular response, as bivalent AAHI-YFV/ZKV vaccination induced levels of antigen-reactive polyfunctional CD4^+^ and IFN-γ^+^ CD8^+^ T cells comparable to those induced by the monovalent vaccines. Finally, we demonstrated that bivalent AAHI-YFV/ZKV immunization, at various doses, conferred complete protective efficacy in both mouse and hamster models of lethal ZIKV and YFV challenge. Taken together, the potent induction of humoral and cellular responses seen following bivalent vaccination highlights the flexible application of the AAHI saRNA-NLC platform against multiple viruses in a multivalent format.

The presence of serum virus-specific nAbs is critical for protection against flavivirus infection.(44,45) We have previously seen strong and durable induction of nAbs following one-or two-dose immunization with NLC-formulated ZIKV saRNA.(16) The nAb GMT induced following monovalent and bivalent vaccination with AAHI-YFV and/or AAHI-ZKV are consistent with prior work, and are above protective PRNT_50_ values for YFV (PRNT_50_ ≥10 in humans) and ZIKV (PRNT_50_ ≥100 in mouse passive transfer studies) observed by other groups.(2,32,38) Moreover, the magnitude of the nAb responses are comparable to other vaccine strategies targeting YFV(32,42,46–52) and ZIKV.(38,39,53–62) The strong dose-dependent nAb titers induced by AAHI-YFV were notable, which required a high (30 µg) dose or boost vaccination to approach the nAb titers induced in mice by YF-17D immunization. Although levels of YFV E protein-binding IgG and durability of nAb titers were equivalent between AAHI-YFV and YF-17D vaccinated mice, these data suggest a discrepancy in the induction of antigen-binding versus virus-neutralizing antibodies with the current AAHI-YFV saRNA construct. For example, when compared against AAHI-ZKV at comparable saRNA doses, AAHI-YFV induced similar levels of E protein-binding IgG but lower nAb titers. Structural and antigenic differences between prM-E proteins, rather than limitations of the bivalent vaccine strategy, may underpin the differences in antigen-binding versus virus-neutralizing antibodies. Future studies could explore further optimization of the YFV prM-E antigen in an attempt to enhance the ratio of nAbs, for example potentially matching the YFV prM-E structure and function to that of more immunogenic ZIKV prM-E variants.(30,63) While induction of serum nAb titers is considered the primary CoP for flavivirus vaccination, T cells also play a crucial role in the immune response to viral infections. Vaccination with YF-17D induces a robust and persistent polyfunctional CD4^+^ and CD8^+^ T cell response with broad antigen specificity, capable of conferring protection against YFV challenge in conjunction with humoral immunity or through control of early viral loads, respectively.(64–71) Various ZIKV vaccination approaches have also been reported to elicit antigen-specific cellular responses.(38,57–61,72,73) The induction of antigen-reactive CD4^+^ and CD8^+^ T cell responses by our saRNA-NLC formulations paired with strong nAb responses support the value of this vaccine platform. Interestingly, while induction of cross-reactive CD4^+^ T cells was minimal, cross-reactive CD8^+^ T cell induction was notable following vaccination with both monovalent flavivirus constructs. This difference may be a consequence of the MHC haplotype in C57BL/6 mice, as other groups have reported CD4^+^ T cell flavivirus cross-reactivity using MHC transgenic mouse models(74,75). The consequence of flavivirus cross-reactive cellular responses remains poorly understood but could potentially confer some protection against heterologous flavivirus infections in the absence of robust pre-existing humoral immunity, or shape subsequent T cell responses, as has been proposed for dengue virus and ZIKV.(76,77) A flavivirus cross-reactive cellular response may partially explain the protection conferred by AAHI-YFV and AAHI-ZKV vaccination against ZIKV and YFV challenge, respectively; however, this possibility requires additional confirmation, such as through passive serum transfer and T cell depletion studies. In sum, the ability of this saRNA-NLC vaccine platform to stimulate strong T cell responses is promising, both for these flaviviral-targeted vaccines as well as vaccines targeting other infectious diseases.

To date, only one YFV vaccine has been approved for commercial use, the live attenuated strain YF-17D. Notably, while the safety profile of YF-17D is generally excellent in immunocompetent populations, and manufacturers stockpile 30-80 million doses/year, viscerotropic effects at the extremes of age, dissemination in the immunocompromised, and supply limitations have left public health officials without a complete set of tools for the prevention of YFV despite this durable and highly immunogenic YFV vaccine.(6) In contrast with YFV, there are currently no approved ZIKV vaccines, in large part due to difficulties in conducting Phase III efficacy trials following the decline in circulating ZIKV.(62) ZIKV still presents pandemic risk in the coming years as carrier mosquito populations continue to expand, and there is a potential risk of mutants escaping existing immunity or current vaccine candidates.(78,79) These unmet medical and public health concerns underscore the need for effective, safe, stable, and rapidly producible YFV and ZIKV vaccines. The potent saRNA-NLC vaccine platform presented here is well suited to address these needs, and thus AAHI-YFV/ZKV is a strong candidate for a flavivirus bivalent vaccine.

While the studies conducted here were carefully designed for scientific rigor and reproducibility, some limitations exist. Groups were sex balanced, and while no significant sex-based differences were observed in any of the immunogenicity measures, larger study groups would be necessary to assess more subtle sex-based differences in vaccine-driven immune responses or efficacy against ZIKV challenge. Additionally, no immunogenicity or efficacy data are presented from primate models, which will be a key future direction for these vaccine candidates.

Previous platform development work has demonstrated the inherent flexibility of saRNA synthesis, ease of saRNA and NLC manufacturing, and the stability of lyophilized NLC-formulated saRNA at room temperature for extended periods, allowing for rapid scale-up and stockpiling of vaccine material for emerging viral variants.(17,80) Taken together, our saRNA-NLC system has the potential to improve global access to flavivirus vaccines, particularly in resource-limited areas, and serve as a platform to develop additional vaccines in multivalent format.

## DATA AVAILABILITY STATEMENT

The raw data supporting the conclusions of this article will be made available by the authors, without undue reservation.

## ETHICS STATEMENT

All mouse work was done under the oversight of the Bloodworks Northwest Research Institute’s (Seattle, WA) Institutional Animal Care and Use Committee (IACUC), protocol #5389-01. All hamster care and husbandry were conducted according to protocols approved by the Utah State University IACUC, protocol #10010.

## AUTHOR CONTRIBUTIONS

CC and EAV conceived the study. MRY, MFJ, AG, JGJ, and EAV supervised the research. AG, JGJ, CC, and EAV administered the project. EAV acquired funding. PB, MRY, MFJ, SB, AEW, DH, JS, JB, NC, PF, JA, and SR contributed to the investigation. PB, MRY, MFJ, DH, and JB contributed to methodology. PB, MRY, MFJ, SB, DH, JS, JB, SR, AG, and EAV performed validation. PB, MRY, DNK, MFJ, SB, JS, and SR curated the data. PB, MRY, DNK, MFJ, SB, JS, and EAV analyzed the data. PB, MRY, and DNK visualized the data. PB, MRY, and DNK wrote the initial manuscript. PB, MRY, DNK, AG, CC, and EAV revised the manuscript. PB, MRY, DNK, and EAV accessed and verified the data. All authors reviewed and approved the final version of the manuscript.

## FUNDING

This project has been funded in whole or in part with Federal funds from the National Institute of Allergy and Infectious Diseases of the National Institutes of Health, under Contract No. 75N93019C00059.

## CONFLICT OF INTERESTS

AG and EAV declare no Competing Non-Financial Interests but the following Competing Financial Interests. AG and EAV are co-inventors on U.S. patent application nos. PCT/US21/40388, “Co-lyophilized RNA and Nanostructured Lipid Carrier,” and 63/144,169, “A thermostable, flexible RNA vaccine delivery platform for pandemic response.” All other authors declare that they have no competing interests.

