## Supplementary Info for "A bivalent self-amplifying RNA vaccine against yellow fever and Zika viruses"

### SUPPLEMENTARY INFORMATION

#### SUPPLEMENTARY METHODS

##### saRNA production

YFV-prM-E saRNA and ZIKV-prM-E saRNA were synthesized at 200 mg scale using an AAHI-optimized in vitro transcription protocol. Briefly, saRNA template plasmids were created by subcloning the YFV and ZIKV prM-E inserts into the plasmid T7-VEEVRep, which contains a 5' UTR, a 3' UTR, and non-structural proteins derived from the attenuated TC83 strain of Venezuelan equine encephalitis virus (VEEV). Flaviviral prM-E sequences were cloned downstream of the sub-genomic promoter, in place of VEEV structural proteins. Generation of saRNA was achieved by T7-mediated polymerization from a restriction-endonuclease linearized DNA template and capping using vaccinia capping enzyme. saRNA was then purified using flow-through mode chromatography with a CaptoCore 700 resin to remove protein components followed by concentration and diafiltration by tangential flow filtration with a 750 kDa cutoff MidiKros-mPES hollow fiber membrane into a 10 mM Tris pH 8 buffer. Each saRNA batch was terminally filtered with a 0.22 µm polyethersulfone filter and stored at -80°C.

##### NLC production

NLC was produced by combination of an oil phase (squalene, sorbitan monostearate, DOTAP (N-[1-(2,3-dioleoyloxy)propyl]-N,N,N-trimethylammonium chloride), and glyceryl trimyristate) with an aqueous phase (10 mM sodium citrate trihydrate buffer and polysorbate 80) in a two-step process. The two phases were separately heated to 70°C in a bath sonicator. Following complete dissolution of the solid components, the oil and aqueous phases were mixed at 7,000 rpm in a high-speed laboratory emulsifier (Silverson Machines) to produce a crude mixture containing micron-sized oil droplets. This mixture was further processed via high-shear homogenization in a M-110P microfluidizer (Microfluidics) at 30,000 psi to produce the final NLC particles. The batch was terminally filtered with a 0.22 µm polyethersulfone filter and stored at 2-8°C.

##### Vaccine complexing and bivalent dose preparation

Monovalent vaccine complexes for *in vivo* injection were created by mixing diluted aqueous RNA 1:1 by volume with diluted NLC in a final formulation background of 10% w/v sucrose, 5 mM sodium citrate. All vaccines were prepared at a nitrogen:phosphate (N:P) ratio of 15 and were isotonic. Each monovalent vaccine was incubated on ice for 30 minutes after mixing to ensure complete complexing. Three different mixing strategies were used to prepare the bivalent groups (Supplementary Table 1). The first strategy delivered the bivalent vaccine by injecting each monovalent vaccine in a separate leg ("split monovalent"). The second strategy ("complex -> mix") mixed the monovalent vaccines together 1:1 by volume after they had completed the 30-minute incubation step. In the third strategy ("mix -> complex"), the two saRNA components were mixed together first and then complexed with NLC as described above. After the study evaluating the three different bivalent dosing strategies, subsequent bivalent vaccines were prepared using the "complex -> mix" strategy. In all studies, vaccines were frozen at -80°C prior to shipment and/or administration.

**Transfection, protein harvest, and western blotting**

12-well tissue culture treated plates were plated with 4 x $10^5$ HEK293T cells (American Type Culture Collection #CRL-11268) in 1 mL of Dulbecco's Modified Eagle Medium (DMEM) + GlutaMAX (Gibco) with 10% (v/v) fetal bovine serum (FBS) and incubated overnight. Vaccine was complexed as above and then diluted to 10 ng/µL. Media were aspirated from the 12-well plates, and 450 µL of serum-free Opti-MEM (Gibco) and 50 µL of vaccine complex was added to each well. After a 4-hour incubation, media were aspirated, and 1 mL of DMEM with 1% (v/v) FBS was added. Plates were incubated at 37°C with 5% $CO_2$ for 5-48 hours, with samples collected throughout the time course. Cell supernatant was collected, followed by cell lysate harvest using RIPA buffer (Thermo Fisher Scientific). YFV and ZIKV E protein expression was measured by western blot of transfected cell supernatants and lysates. Tris-glycine gels were run and transferred to a polyvinylidene difluoride membrane using an iBlot 2 (Invitrogen) system. Membranes were incubated in primary antibody against YFV (clone I95F, Invitrogen #MA1-7398) or ZIKV (polyclonal, GeneTex #GTX133314) diluted in blocking buffer, then rinsed and incubated in secondary antibody (Invitrogen #A10685 or GeneTex #GTX213110-01) prior to rinsing and development using SuperSignal West Pico PLUS Chemiluminescent Substrate (ThermoFisher) and imaging using a ChemiDoc MP imaging system (Bio-Rad).

**Animal studies**

*Mouse studies*

Mice were housed 3-5 animals per cage with irradiated bedding in individually ventilated cages. During the ZIKV challenge study, some male mice were separated due to fighting and subsequently housed individually for the remainder of the study. Sterile water gel packs were given to all newly arrived animals during the acclimation period, and animals had access to fresh potable water ad libitum. Cages were changed every 2 weeks inside a laminar-flow cage change station, enrichment bedding, and engagement items were changed the same frequency, and spot checks were conducted on non-change weeks. Complexed saRNA-NLC vaccines were thawed on wet ice at the time of immunization for prime (Day 0) or boost (Day 28) immunizations and took place within a 1-hour time window not more than 4 hours after vaccine thawing. Serum samples were taken by retro-orbital bleed (up to 100 µL for survival samples) under isoflurane sedation to mitigate animal pain and distress or cardiac bleed (for terminal samples). Mice were humanely euthanized by $CO_2$ overdose followed by cervical dislocation at terminal harvest for serum and spleen samples. Criteria for euthanasia included lack of mobility, lethargy, hunched back without resolution, weight loss greater than 20% relative to entire study population or challenge day 0, and end of viral challenge period.

*Hamster studies*

Animals were weighed prior to the start of the study and weekly thereafter. If morbidity was observed or weight loss suspected outside of the challenge phase, hamsters were weighed more frequently up to a maximum of daily and compared to the entire study population. Complexed saRNA-NLC vaccines were thawed on wet ice at the time of immunization for prime (Day 0) or boost (Day 28) immunizations and took place within a 1-hour time window not more than 4 hours after vaccine thawing. Animals were observed daily for 3 days after each immunization. Serum samples were obtained as approved by ocular sinus bleed (up to 150 µL for survival samples) or cardiac bleed (for terminal samples). Hamsters were immunized through intramuscular injection bilaterally in the rear quadriceps muscles (50 µL/leg, 100 µL total). Criteria for euthanasia included lack of mobility, inability to right self or access food and water, and weight loss greater than 20%.

**ELISA**

Plates (384-well high-binding plates, Corning) were coated with 25 µl/well of 1 µg/mL of recombinant YFV envelope protein (Meridian Bioscience #R01709) or recombinant ZIKV envelope protein (Meridian Bioscience #R01635) in phosphate-buffered saline (PBS) and incubated overnight at 4°C. The coating solution was removed, and blocking buffer (2% dry milk, 0.05% Tween 20, and 1% goat serum) was applied for at least 1 hour. On a separate, low-binding plate, each serum sample was diluted 1:40 and then serially 1:2 to create a 14-point dilution curve for each sample. Naïve mouse serum, used as a negative control, was diluted identically. A pan-flavivirus envelope protein-binding monoclonal antibody (mAb) (hybridoma clone 4G2; Novus Biologicals #NBP2-52709) was used as a positive control at a known starting concentration of 3.2 ng/µL followed by serial 1:2 dilutions similarly to each sample and negative control. Protein-coated and blocked assay plates were washed, and serially diluted samples and controls were then transferred onto the coated plates followed by a 1-hour incubation. Plates were then washed, and protein-bound antibodies were detected using an anti-mouse IgG (Fc specific)-alkaline phosphatase antibody (Sigma-Aldrich #A2429) at a 1:4000 dilution in blocking buffer. Plates were washed and then developed using phosphatase substrate tablets (Sigma-Aldrich #S0942) dissolved in diethanolamine substrate buffer (Fisher Scientific #PI34064) at a final concentration of 1 mg/mL. After a 30-minute development, plates were read spectrophotometrically at 405 nm. A 4-point logistic (4PL) curve was used to fit the antibody standard titration curve. Sample concentrations were interpolated off the linear region of each sample dilution curve using the standard curve for absolute quantification of antibody titers.

**Plaque reduction neutralization test**

Vero cells (ATCC #CCL-81) were plated in 6-well plates at 5 x $10^5$ cells per well in DMEM (Gibco) with heat-inactivated 10% FBS (Cytiva). Serum samples were diluted 1:10 in DMEM with 1% FBS, then serially diluted 1:2. A 100 PFU/115 µL dose of YF-17D virus or ZIKV FSS13025 was added to each serum dilution. Mixtures were incubated at 37°C for 1 hour and then transferred to the 6-well plates. Plates were incubated at 37°C for 1 hour, followed by media removal and overlay with DMEM with 1% FBS and 0.3% melted agarose. Overlays were allowed to solidify at room temperature after which the plates were incubated at 37°C for 3 days (ZIKV) or 5 days (YFV). Plates were fixed with 10% neutral buffered formalin (Fisher Scientific), and plaques were visualized with Crystal violet (BD). Plaque reduction neutralization test (PRNT) titers were calculated as the last dilution with ≥ 50% reduction of plaques compared to the virus–DMEM only control ($PRNT_{50}$).

**Flow cytometry**

Spleens were collected 28 days post-prime or post-boost and processed by mechanical dissociation to achieve a single-cell suspension. Briefly, spleens were dissociated in RPMI medium through a 70 µm cell strainer by maceration using a syringe plunger flange. Homogenized samples were centrifuged at 400 x g for 5 minutes, supernatant was discarded, and cells were treated with RBC lysis buffer. RPMI medium was added to quench the lysis, and tubes were centrifuged at 400 x g for 10 minutes, after which the supernatant was discarded. Cell pellets were resuspended in 1 mL of RPMI medium and then filtered through a 96-deep-well filter plate (AcroPrep, Cytiva) by centrifugation at 400 x g for 5 minutes. Splenocytes were resuspended in RPMI medium with 10% FBS, seeded in 96-well round-bottom plates at 1-2 x $10^6$ cells/well, and stimulated at 37°C with 5% $CO_2$ for 6 hours. Stimulations consisted of media (RPMI, 10% FBS, 50 µM 2-mercaptoethanol, α-CD28 [BD # BDB553294], and brefeldin A [BioLegend]) with YFV or

ZIKV prM-E protein peptide pools (15-mers with 11 aa overlap) resuspended in dimethyl sulphoxide (DMSO) (1 µg/mL, GenScript); DMSO serving as a negative control; or Cell Activation Cocktail without brefeldin A (BioLegend) serving as a positive control. After stimulation, plates were centrifuged at 400 x g for 3 minutes, the supernatants were removed by pipetting, and cells were washed with 1X PBS. Splenocytes were stained for viability with Zombie Green (BioLegend), and Fc receptors were blocked with α-CD16/CD32 (Invitrogen). Cells were surface stained for CD4 (PerCP-Cy5.5, BioLegend #100434 or 100540), CD8α (BV510, BD Biosciences #563068), and CD44 (APC-Cy7, BioLegend #103028) in staining buffer (1X PBS, 0.5% BSA, and 0.1% sodium azide) for 20 minutes on ice. Cells were permeabilized using the BD Cytofix/Cytoperm Kit (BD Biosciences) and stained intracellularly for TNF-α (BV421, BioLegend #506328), IL-2 (PE-Cy5, BioLegend #503824), IFN-γ (PE-Cy7, BD Biosciences #557649), IL-5 (PE, BioLegend #504304), and IL-17a (AF700, BD Biosciences #560820). Samples were run on a Fortessa flow cytometer (BD) and analyzed using FlowJo software (BD Biosciences).

**Cells and viruses**

Vero cells (ATCC #CCL-81 and CRL-1587) were maintained at 37°C with 5% $CO_2$ in DMEM containing 10% heat-inactivated FBS (Cytiva #SH30396.03HI), sodium pyruvate (1 mM), and penicillin-streptomycin (100 U/mL each, Gibco #15140122). Cell lines were pre-screened for mycoplasma contamination.

Passage 11 (P11) of the Jiménez strain was used for YFV challenge. The working stock was prepared as previously described(1,2) through a single passage in hamsters. Three female hamsters were infected intraperitoneally with P10 (obtained from the World Reference Center for Emerging Viruses and Arboviruses, University of Texas Medical Branch, Galveston, TX), and livers were collected 3 days post-infection and homogenized in a 2X volume of sterile PBS. This challenge lot typically results in lethality of 80%, which is dose independent (i.e., higher challenge doses do not cause greater mortality). For $PRNT_{50}$, Yellow Fever Virus, YF-17D (NR-116) was obtained through BEI Resources (NIAID, NIH), and was passaged once in Vero cells. These viral stocks were titrated by plaque assay on Vero cell monolayer cultures prior to storage at -80°C.

A P2 stock of the MA-Zika-Dakar strain (GenBank ID: MG758786.1) was used for ZIKV challenge. The working stock was generated and propagated as previously described.(3–5) The ZIKV FSS13025 strain was obtained as a generous gift from Michael Gale (University of Washington, Seattle, WA) and passaged once in the *Aedes albopictus* clone C6/36 prior to storage at -80°C and use in PRNT assays.

**Viral titer measurement**

Peripheral serum drawn from infected hamsters 4 days post-infection and stored at -80°C until assessment for viremia by infectious virus assay.(6) Virus was titrated in 96-well plates containing Vero 76 cells (ATCC #CRL-1587). Ten-fold serial dilutions of serum samples were incubated with confluent monolayers at 37°C. Plates were read when cytopathic effect was apparent around 7 days after plating. All samples were assayed in triplicate.

**Alanine aminotransferase assay**

A colorimetric assay for the detection of serum alanine aminotransferase (ALT) (Teco Diagnostics) was used to determine the level of ALT in the serum of infected hamsters collected 6 dpi. The manufacturer’s protocol was modified to use 10-fold less sample volume and reagents in a 96-well plate.

**Replicates**

Replication of data was conducted whenever possible. All *in vitro* and *in vivo* studies were conducted with a minimum of biological triplicates, as indicated. ELISAs were conducted with technical duplicates for each study sample. Splenocytes samples were split into three aliquots, which were each stimulated and stained independently as technical triplicates prior to pooling and enumeration of antigen-specific T cells by flow cytometry. Bridging groups were used between independent animal studies to provide replication of key study groups across multiple studies offset by months.

**Sample size estimation**

We have previously determined that 10 mice per group, 5 male and 5 female, provides statistical power of at least 90% with an alpha (*p*-value) of 0.05 to detect a statistically significant difference in ELISA and $PRNT_{50}$ antibody titers between vaccinated groups with two-fold differences in mean antibody titers with expected statistical variance. Sample sizes of 6 and 10 animals per study group were previously found to provide sufficient statistical power to detect differences in vaccine immunogenicity and efficacy for ZIKV challenge studies. Therefore, 6 or 10 mice, for immunogenicity or efficacy, respectively, were used for each group, for a total of 5-6 or 12 groups (50-60 or 72-120 animals) per study. Sample sizes of 10 hamsters per group, 5 male and 5 female, were used and were previously found to provide sufficient statistical power to detect differences in vaccine efficacy for YFV challenge studies. Therefore, 10 hamsters were used for each group, for a total of 14 groups (140 animals) per study.

**Randomization**

Mice were randomized between groups within sex. Hamsters were randomly and blindly distributed into respective study groups within sex.

**Blinding**

Investigators were not blinded to the study groups during data collection and analysis. Quantitative, unbiased assays were developed with highly regulated SOPs to minimize data biases.

**Inclusion and exclusion criteria**

No data were excluded from any plots or analyses, except for flow cytometry where samples with insufficient threshold events (<50,000) collected per well were excluded.


**SUPPLEMENTARY TABLES**

**Supplementary Table 1. Experimental groups for bivalent dosing study.**

| **Animal group** | **RNA replicon** | **RNA concentration (ng/µL)** | **NLC concentration (% DOTAP)** | **RNA dose in mice (µg)** | **Injection site** |
|---|---|---|---|---|---|
| **1** | SEAP | 200 | 0.6 | 10 | 1x rear leg |
| **2** | YFV | 200 | 0.6 | 10 | 1x rear leg |
| **3** | ZIKV | 200 | 0.6 | 10 | 1x rear leg |
| **4A** | YFV | 200 | 0.6 | 20 (10 per construct) | Monovalent injections, separate rear legs (Split Monovalent) |
| **4B** | ZIKV | 200 | 0.6 | | |
| **5** | YFV-ZIKV | 400 (200 per construct) | 1.2 | 20 | Bivalent injection, mixed *post-complexing*, 1x rear leg (Complex -> Mix) |
| **6** | YFV-ZIKV | 400 (200 per construct) | 1.2 | 20 | Bivalent injection, mixed *pre-complexing*, 1x rear leg (Mix -> Complex) |

**Supplementary Table 2. Effect of AAHI-YFV and AAHI-ZKV vaccination in a mouse model of Zika virus challenge.**

| | | | | | |
|---|---|---|---|---|---|
| *Animals*: C57BL/6J | N/A | N/A | N/A | *Duration of experiment*: 72 days | N/A |
| *Virus, route*: $10^5$ PFU MA-Zika Dakar, IP | N/A | N/A | N/A | *Treatment volume:* 100 µL IM | N/A |
| **Treatment** | **Schedule** | **Sex** | **Alive/Total** | **MDD[1] ± SD** | **Mean wt. change[2] (g) ± SD** |
| 10 µg SEAP | d0 | M; F | 0/5; 4/5 | 8.6 ± 0.8; 12.8 ± 2.4 | -2.04 ± 0.467; -1.180 ± 0.396 |
| 10 µg AAHI-YFV | d0 | M; F | 4/5; 3/5 | 13.6 ± 0.8; 11.4 ± 3.2 | 0.02 ± 0.286****; -0.820 ± 1.119 |
| 10 µg AAHI-ZKV | d0 | M; F | 5/5; 5/5 | 14 ± 0.0**; 14 ± 0.0 | 0.220 ± 0.559****; 0.14 ± 0.167*** |
| 10/10 µg AAHI-YFV/ZKV | d0 | M; F | 5/5; 5/5 | 14 ± 0.0**; 14 ± 0.0 | 0.260 ± 0.483****; 0.4 ± 0.212**** |
| 5/5 µg AAHI-YFV/ZKV | d0 | M; F | 5/5; 5/5 | 14 ± 0.0**; 14 ± 0.0 | 0.04 ± 0.611****; 0.52 ± 0.239**** |
| 1/1 µg AAHI-YFV/ZKV | d0 | M; F | 5/5; 5/5 | 14 ± 0.0**; 14 ± 0.0 | 0.1 ± 0.224****; 0.58 ± 0.192**** |
| 10 µg SEAP | d0, d28 | M; F | 0/5; 4/5 | 8.8 ± 0.4; 13.4 ± 1.2 | -2.04 ± 0.434; -1.22 ± 0.409 |
| 10 µg AAHI-YFV | d0, d28 | M; F | 3/5; 5/5 | 12.2 ± 2.2; 14 ± 0.0 | -0.96 ± 1.099; -0.18 ± 0.873 |
| 10 µg AAHI-ZKV | d0, d28 | M; F | 5/5; 5/5 | 14 ± 0.0**; 14 ± 0.0 | -0.16 ± 0.873***; -0.16 ± 0.297 |
| 10/10 µg AAHI-YFV/ZKV | d0, d28 | M; F | 5/5; 5/5 | 14 ± 0.0**; 14 ± 0.0 | -0.52 ± 0.396**; 0.44 ± 0.770*** |
| 5/5 µg AAHI-YFV/ZKV | d0, d28 | M; F | 5/5; 5/5 | 14 ± 0.0**; 14 ± 0.0 | -0.12 ± 0.487***; 0.84 ± 0.472**** |
| 1/1 µg AAHI-YFV/ZKV | d0, d28 | M; F | 5/5; 5/5 | 14 ± 0.0**; 14 ± 0.0 | 0.14 ± 0.219****; 0.34 ± 0.336** |

[1]Mean day of death.
[2]Difference between weight on 3 and 6 days post-virus challenge representing maximal weight change within this study.
*p<0.05, **p<0.01, ***p<0.001, ****p<0.0001 as compared to SEAP-expressing vector control immunization.

**Supplementary Table 3. Effect of AAHI-YFV and AAHI-ZKV vaccination in a hamster model of yellow fever virus challenge.**

| | | | | | | |
|---|---|---|---|---|---|---|
| *Animals*: Syrian golden hamsters | N/A | N/A | *Duration of experiment*: 80 days | N/A | N/A | N/A |
| *Virus, route*: 200 $CCID_{50}$ YFV Jimenez, IP | N/A | N/A | *Treatment volume:* 100 µL IM | N/A | N/A | N/A |
| **Treatment** | **Schedule** | **Alive/ Total** | **MDD[1] ± SD** | **Mean wt. change[2] (g) ± SD** | **Viremia[3]** | **ALT[4]** |
| 10 µg SEAP | d0 | 2/10 | 7.6 ± 0.9 | -13 ± 7.0 | 6.5 ± 1.2 | 218 ± 68 |
| 10 µg AAHI-YFV | d0 | 10/10 | >21.0 ± 0.0*** | 2.4 ± 2.2*** | 1.7 ± 0.0*** | 88 ± 20*** |
| 10 µg AAHI-ZKV | d0 | 6/10 | 8.3 ± 1.0* | -8.0 ± 8.7 | 5.7 ± 2.1 | 138 ± 36*** |
| 10/10 µg AAHI-YFV/ZKV | d0 | 10/10 | >21.0 ± 0.0*** | 1.9 ± 2.6*** | 1.7 ± 0.0*** | 88 ± 16*** |
| 5/5 µg AAHI-YFV/ZKV | d0 | 10/10 | >21.0 ± 0.0*** | 2.7 ± 3.0*** | 1.7 ± 0.0*** | 87 ± 16*** |
| 1/1 µg AAHI-YFV/ZKV | d0 | 9/10 | 7.0 ± 0.0** | 1.0 ± 6.2*** | 2.6 ± 1.7**** | 91 ± 39*** |
| $10^4$ PFU YFV-17D | d0 | 10/10 | >21.0 ± 0.0*** | 2.6 ± 3.3*** | 1.8 ± 0.2*** | 91 ± 52*** |
| 10 µg SEAP | d0, d28 | 1/10 | 8.4 ± 0.9 | -6.8 ± 5.4 | 6.2 ± 1.3 | 178 ± 74 |
| 10 µg AAHI-YFV | d0, d28 | 10/10 | >21.0 ± 0.0*** | 2.0 ± 2.3*** | 1.7 ± 0.0*** | 85 ± 21*** |
| 10 µg AAHI-ZKV | d0, d28 | 10/10 | >21.0 ± 0.0*** | 0.7 ± 4.6** | 3.0 ± 1.8*** | 113 ± 39*** |
| 10/10 µg AAHI-YFV/ZKV | d0, d28 | 9/10 | 7.0 ± 0.0*** | 0.8 ± 2.7** | 1.7 ± 0.0*** | 81 ± 9.2*** |
| 5/5 µg AAHI-YFV/ZKV | d0, d28 | 10/10 | >21.0 ± 0.0*** | 2.4 ± 3.2*** | 1.7 ± 0.0*** | 85 ± 15*** |
| 1/1 µg AAHI-YFV/ZKV | d0, d28 | 10/10 | >21.0 ± 0.0*** | 2.6 ± 3.2*** | 1.7 ± 0.0*** | 79 ± 7.0*** |
| $10^4$ PFU YFV-17D | d0, d28 | 8/8 | >21.0 ± 0.0*** | 2.4 ± 2.0*** | 1.7 ± 0.0*** | 79 ± 13*** |
| Normal Controls | NA | 3/5 | 14 ± 0.0 | 0.0 ± 1.9*** | 1.7 ± 0.0*** | 77 ± 7.0*** |

[1]Mean day of death.
[2]Difference between weight on 3 and 6 days post-virus challenge representing maximal weight change within this study.
[3]Serum was collected 4 dpi. Values are mean virus titer ($CCID_{50}$/mL) ± SD.
[4]Serum was collected 6 dpi. Values are mean IU/L ± SD.
*p<0.05, **p<0.01, ***p<0.001, ****p<0.0001 as compared to SEAP-expressing vector control immunization.

**SUPPLEMENTARY FIGURES**

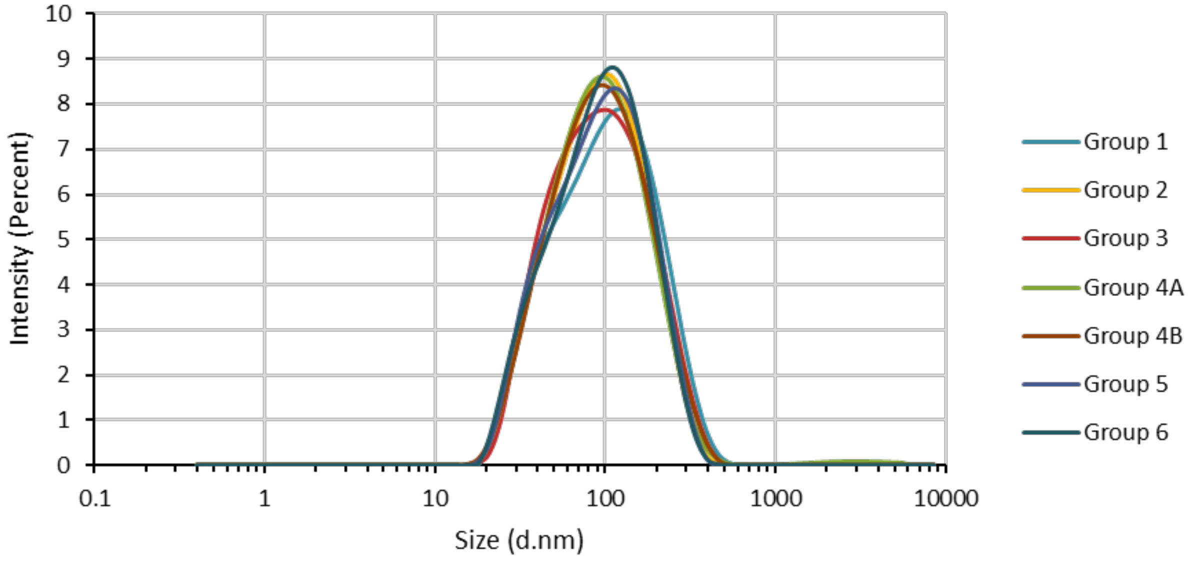


**Supplementary Figure 1. Intensity-based particle size distribution of saRNA-NLC vaccine complexes as measured by dynamic light scattering.** Related to Figures 5 and 6. Intensity-weighted Z-average diameter was determined for each formulation and averaged from three measurements per formulation. Groups defined in Supplementary Table 1.

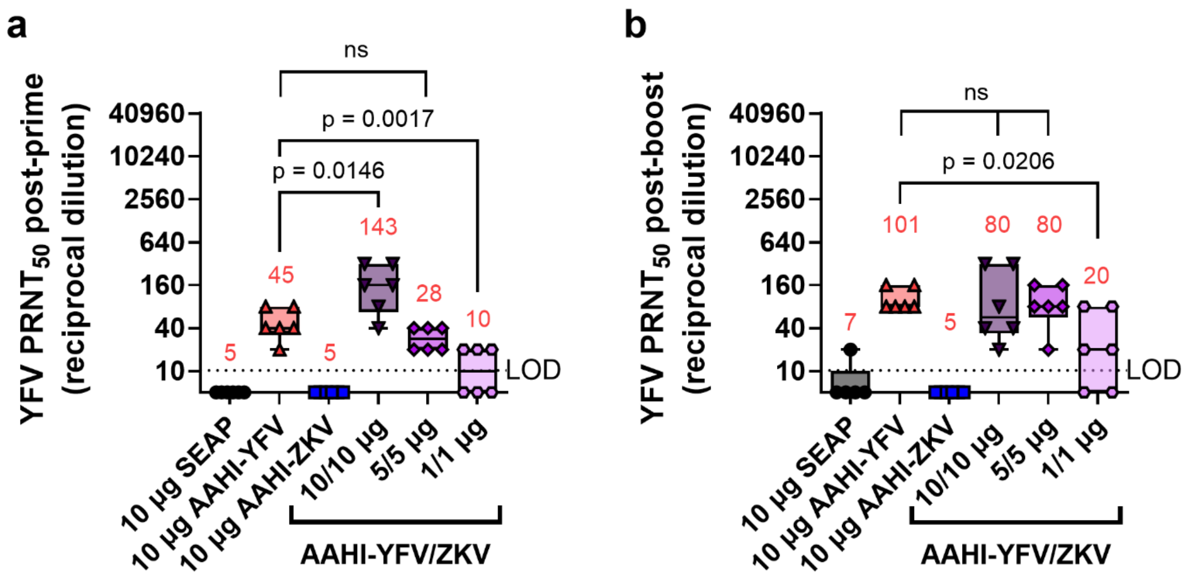


**Supplementary Figure 2. Mouse serum YFV neutralizing titer after monovalent and bivalent saRNA-NLC vaccination.** Related to Figure 7. Serum YFV neutralizing antibody titer 28 days (**a**) post-prime and (**b**) post-boost. Statistical analysis was conducted on $\log_{10}$ transformed data using one-way ANOVA with Dunnett's correction for multiple comparisons. ns = non-significant ($p > 0.05$). Black dotted line shows the limit of detection (LOD) for the assay. Results are from a single independent experiment; $n = 6$ mice per group, 3 male and 3 female. Box plots show median and IQR ± min/max value. Red numerical values represent group $PRNT_{50}$ GMT.

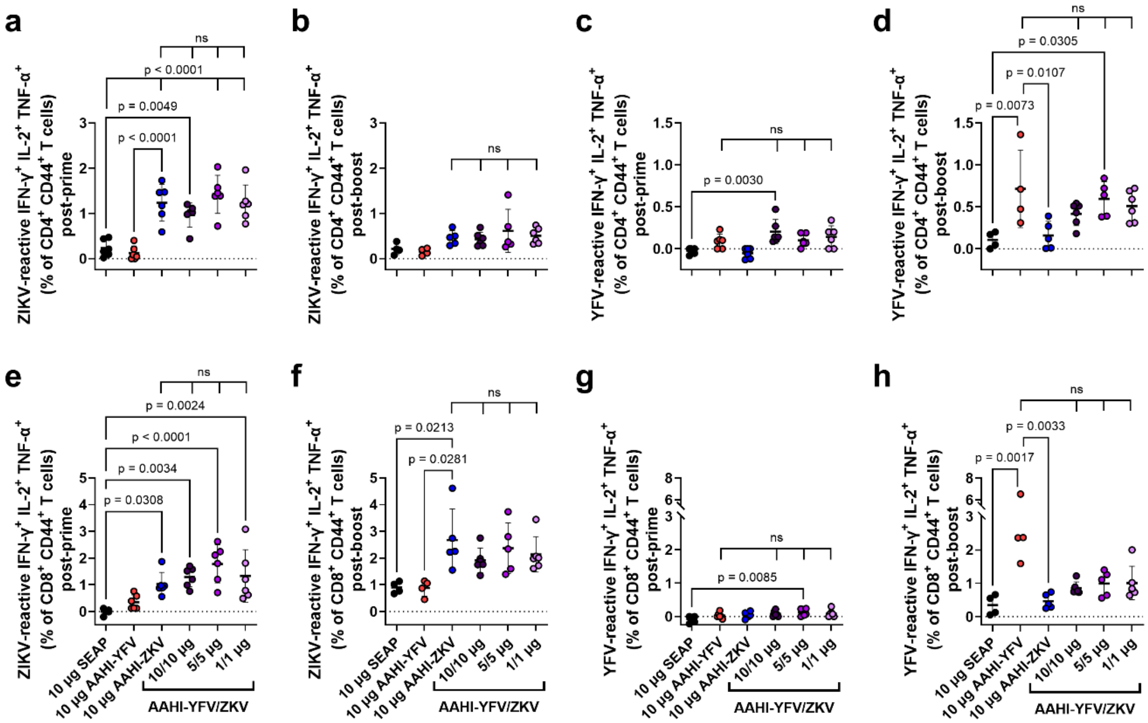


**Supplementary Figure 3. Mouse splenic YFV and ZIKV antigen-reactive T cells after monovalent and bivalent saRNA-NLC vaccination.** Related to Figure 7. Splenic antigen-reactive polyfunctional [IFN-$\gamma^+$ IL-$2^+$ TNF$\alpha^+$] (**a-d**) $CD4^+$ T cells and (**e-h**) $CD8^+$ T cells 28 days (**a, c, e, g**) post-prime and (**b, d, g, h**) post-boost. (**a, b, e, f**) ZIKV-reactive and (**c, d, g, h**) YFV-reactive T cells. Statistics were measured using one-way ANOVA with Šidák's correction (**a-e, g**) or Kruskal-Wallis with Dunn's correction (**f, h**) for multiple comparisons. Comparison were made against 10 µg SEAP-expressing vector control and monovalent AAHI-YFV or AAHI-ZKV. ns= non-significant ($p > 0.05$). Results are from a single independent experiment; $n = 6$ mice per group, 3 male and 3 female. Scatter plots show mean ± SD.

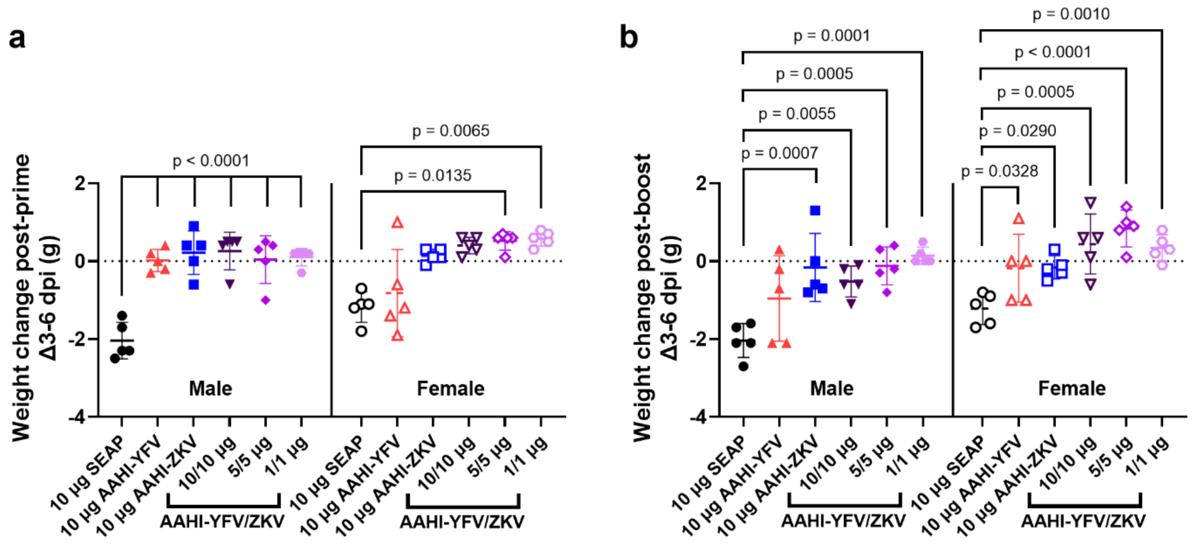


**Supplementary Figure 4. Mouse post-ZIKV challenge morbidity.** Related to Figure 7. (**a**) Post-prime and (**b**) post-boost bodyweight change 3-6 dpi. Statistics were measured using one-way ANOVA with Dunnett's correction (**a**, males; **b**, males and females) or Kruskal-Wallis with Dunn's correction (**a**, females) for multiple comparisons against 10 µg SEAP-expressing vector control. Results are from a single independent experiment; *n* = 10 mice per group, 5 male and 5 female. Scatter plots show mean ± SD.

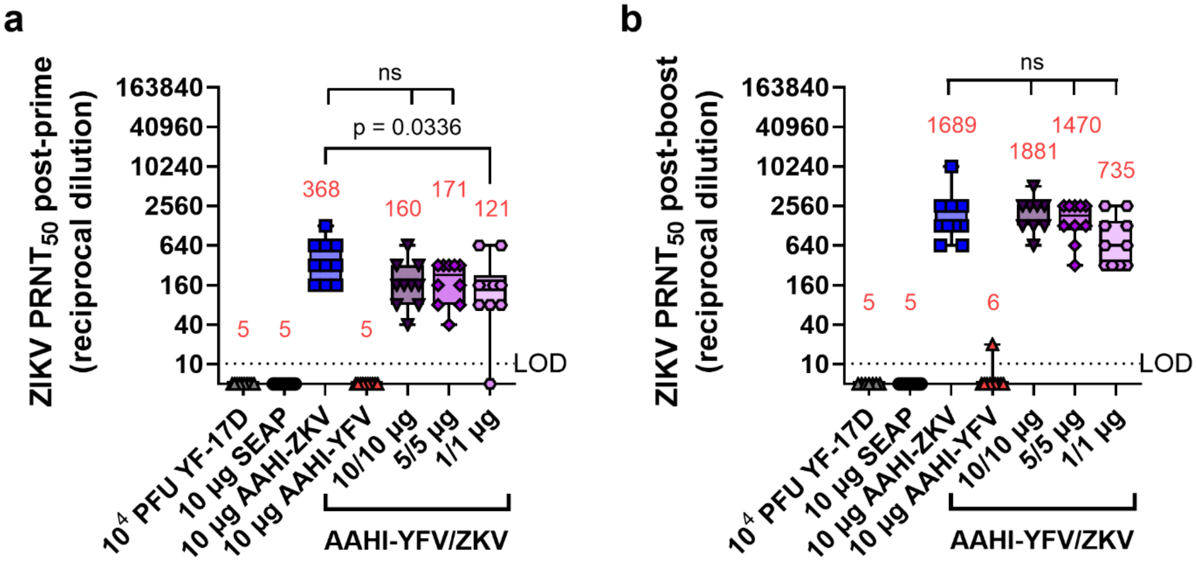


**Supplementary Figure 5. Hamster serum ZIKV neutralizing titer after monovalent and bivalent saRNA-NLC vaccination.** Related to Figure 8. Serum ZIKV neutralizing antibody titer 28 days (**a**) post-prime and (**b**) post-boost. Statistical analysis was conducted on $log_{10}$ transformed data using one-way ANOVA with Dunnett's correction for multiple comparisons. ns = non-significant ($p > 0.05$). Black dotted line shows the limit of detection (LOD) for the assay. Results are from a single independent experiment; $n = 10$ hamsters per group, 5 male and 5 female. Box plots show median and IQR ± min/max value. Red numerical values represent group $PRNT_{50}$ GMT.

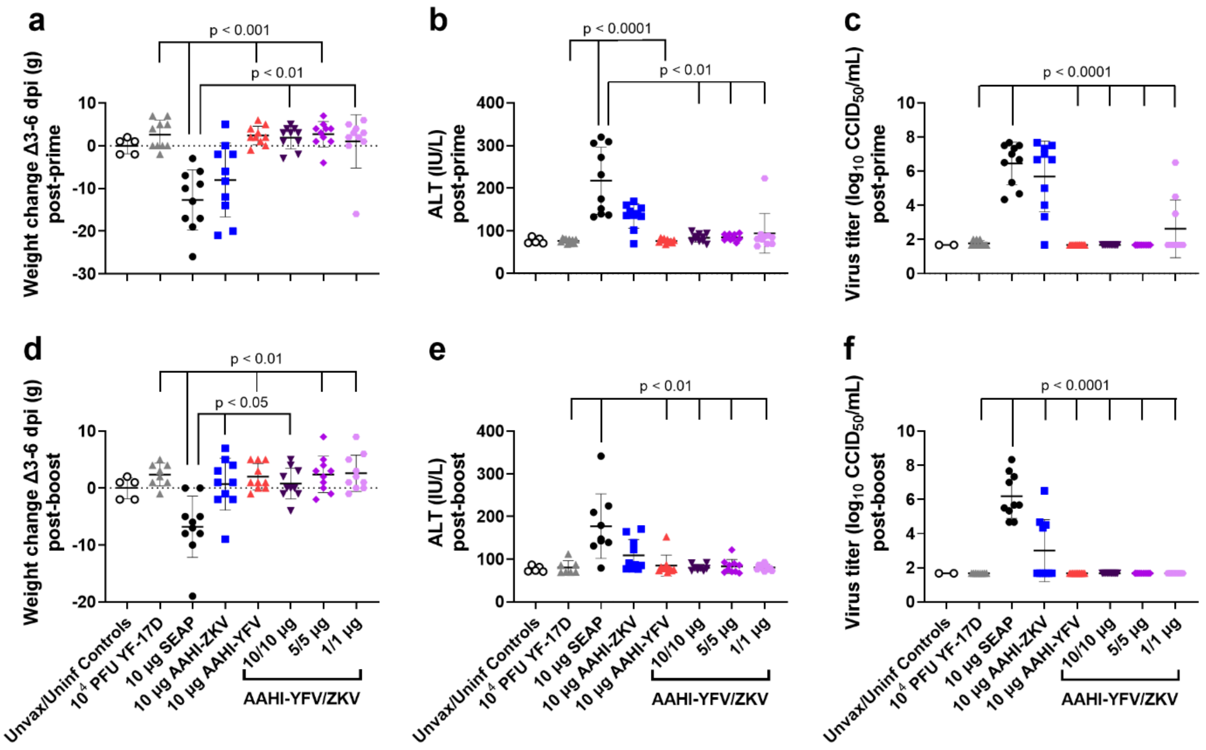


**Supplementary Figure 6. Hamster post-YFV challenge morbidity and viremia.** Related to Figure 8. (**a**) Post-prime and (**d**) post-boost bodyweight change 3-6 dpi. (**b**) Post-prime and (**e**) post-boost serum ALT 4 dpi. (**c**) Post-prime and (**f**) post-boost viremia 4 dpi. Statistics were measured using Kruskal-Wallis with Dunn's correction (**a, b, d, e**) or on $log_{10}$ transformed data using one-way ANOVA with Dunnett's correction (**c, f**) for multiple comparisons against 10 µg SEAP-expressing vector control. *n* = 10 hamsters per vaccination group, 5 male and 5 female, or *n* = 2-5 female hamsters for unvaccinated/uninfected (Unvax/Uninf) controls.
